# An experimental approach to understanding plastic leachate impacts on marine microorganisms

**DOI:** 10.1101/2025.01.30.635782

**Authors:** S. D. M. Maday, K.M. Handley, G. Northcott, J. M. Kingsbury, D. Smith, O. Pantos, G. Lear

## Abstract

Plastics in the world’s oceans are exposed to diverse environmental stressors that fragment them, accelerating the leaching of associated additives. The impact of potentially toxic plastic degradation products and additives on marine microorganisms remains poorly understood. We assessed the impact of plastic leachate on marine microbial communities *in vitro* by exposure to one of four plastic leachates (from linear low-density polyethylene [LLPDE], polyamide-6 [or polycaprolactam; PA6], polyethylene terephthalate [PET] and polylactic acid [PLA]), prepared by immersing plastics in artificial seawater salts broth for three months at 80 °C. Microbial communities were then exposed to different leachates, noting that lower concentrations of plastic additives leached from the more inert plastic types (LLDPE and PET), as determined by GC-MS. PLA-leachate exposed communities differed significantly in composition from other plastic-leachate-exposed communities (PERMANOVA, *P*=0.001) as assessed by 16S rRNA gene and ITS region amplicon sequencing. Communities exposed to PLA-leachate contained a higher proportion of Proteobacteria, specifically *Halomonas* spp., previously reported to degrade LDPE and common plasticisers. Greater relative abundances of Psathyrellaceae fungi also distinguished these communities from those exposed to other plastic leachates. However, despite significant differences in the structure of communities exposed to PLA-leachate, we found no difference in the relative abundances of gene transcripts associated with plastic degradation. While biodegradable plastics such as PLA may persist for shorter times in the environment than more inert plastics, our study indicates greater consequences for marine microbial communities.

## 1. Introduction

The transport and transfer of toxic chemicals by plastics can impact vital marine microbial processes, including rates and extents of primary production (Gregory, 2009) and carbon cycling (Roman and McCarthy, 2010). Many plastics contain a combination of additives - chemicals included during manufacture to improve the polymer’s functional attributes. Functional additives include stabilisers such as benzophenones and benzotriazoles to suppress UV degradation and oxidation (Dintcheva et al., 2007), fillers such as aluminium hydroxide, used as fire retardants (Shrivastava 2018), and plasticisers such as phthalates and adipates, included to enhance plastic malleability. Polymers also contain quantities of substances unintentionally added, such as residual monomers from manufacturing or weathering (Bridson et al., 2023). Most of these additives and unintentionally added substances are not covalently bound to the polymer matrix and will leach out into the environment over time (Hahladakis et al., 2018; Stringer and Johnston, 2001).

Few studies have explored the effects of complex plastic leachates and degradation products on microbial community composition and functioning (Tetu et al., 2020). However, heterotrophic microbes are known to use dissolved organic carbon (DOC) released from plastics. Romera-Castillo et al. (2018) noted an increase in microbial abundance in the presence of low- and high-density polyethylene (LDPE and HDPE) plastic in the dark, compared to light conditions, suggesting that free radicals produced via photodegradation, might negatively impact microbial growth. This highlights the complexity of the problem - plastic degradation mechanisms may both support and hamper the growth and activity of different marine microorganisms. By exploring the effect of recalcitrant homochain polymer plastic compositions (containing only carbon in the polymer backbone) and their more biodegradable heterochain polymer counterparts on marine microbial communities, we may better understand the consequences for marine microbial health of using and improperly disposing of these different plastic types.

To understand the impacts of plastic leachates on marine microorganisms, specifically from those plastics with heteroatoms, we produced leachates from commonly used plastics (formulations of polyamide-6 [PA6], polyethylene terephthalate [PET] and polylactic acid [PLA], as well as homochain linear low-density polyethylene [LLDPE],) via thermal weathering. We analysed the leachate constituents using non-targeted gas-chromatography mass spectrophotometry (GCMS) to determine the diversity of chemicals released. Next, we exposed an *in vitro* marine plastisphere-derived microbial community to these leachate solutions. Over two weeks, changes in microbial abundance and viability were monitored, along with changes in bacterial and fungal community composition. Further, we assessed whether organisms and gene transcripts associated with plastic degradation increased in abundance following exposure to various plastic leachate solutions.

## 2. Material and Methods

### 2.1 Plastic leachate production

Virgin plastic pellets of four types, supplied from Scion^TM^, New Zealand (polylactic acid [PLA], polyethylene terephthalate [PET], nylon 6 [also known as polyamide-6 or PA6] and linear low-density polyethylene [LLDPE]; Table S1), were individually ground into powders using a Thomas Willey® Mini Mill 3383-L40 (Thomas Scientific, Swedesboro, NJ, U.S.A). Particle sizes were estimated using a Mastersizer 3000 (Malvern Panalytical U.K.), set to eight size ranges: <0.01 µm, 0.01 – 0.02 µm, 0.02 - 0.2 µm, 0.2 – 40 µm, 40 – 200 µm, 200 – 500 µm, 500 – 1000 µm and 1000 – 2000 µm. Most powder fragments ranged from 0.2 µm to 500 µm, except for LLDPE, which had 35% of particles above 1000 µm (Figure S1). All plastics underwent thermo-gravimetric analysis (TGA) to measure the inorganic content (Supplementary Materials S1). TGA indicated that inorganic matter concentrations for all polymer types were below the detection limit of ± 0.1% of the original mass of the samples. Additive contents were all presumed to be ≤ 0.5% (Table S1).

Plastic powders were suspended in sterile artificial seawater salts (ASWS) broth (Supplementary Materials S2) to mimic the basal chemical constituents within the marine environment. Four 250 ml replicates of each plastic were thermally weathered for three months in the dark at 80°C, along with a set of control samples (ASWS lacking ground plastics) before being filtered (0.2 µm) to remove smaller particles, including microbial cells, and adjusted to ∼pH 7. Here, we use the term ‘leachate’ to describe all materials < 0.2 µm lost from the parent material, acknowledging that others more frequently describe plastic leachate as the “desorption of chemicals into the surrounding environment” (Delaeter et al., 2022; Gardon et al., 2020), often ignoring the fate of microparticles within the solution. Before exposure to microbial communities, each plastic leachate was tested for microbial contamination (Supplementary Materials S3).

### 2.2. Marine biofilm community sampling

Plastisphere biofilm material was removed from LLDPE, PA6, PET and PLA paddles (Figure S2) installed in the water of Auckland’s Viaduct Basin for two weeks, as described in the Supplementary Materials (S4). Samples were stored in 15% glycerol at −80°C until required.

### 2.3. Exposing marine plastic-associated microbial communities to plastic leachates

#### 2.3.1 Pooling, preconditioning and characterisation of a basal marine-derived plastic-associated microbial starter community

To provide a single starting community, loops full of inoculum, collected from the thawed biofilm stocks of each plastic type, were composited to inoculate 2 L of Difco Marine Broth 2216 which contains carbon in the form of peptone and yeast extract (see manufacturer notes). Difco Marine Broth 2216 was chosen for its propensity to promote the growth of heterotrophic marine bacteria. The inoculated medium was shaken at room temperature (18 – 22°C) for seven days, together with an uninoculated negative control (this is referred to as the preconditioning period, which allowed time for the community to become somewhat used to incubation under laboratory conditions). Prior to leachate exposure, 20 ml of preconditioned culture were aliquoted, in triplicate, for RNA, DNA and GC-MS analysis (i.e., a total of nine aliquots). Aliquots for RNA and GC-MS analyses were snap-frozen using liquid nitrogen and stored at −80°C, while aliquots for DNA analysis were stored at − 20°C until needed. During the preconditioning period (every 24 h), cell counts were estimated via optical density measures (OD_600nm_) and use of a LIVE/DEAD *Bac*Light Bacterial Viability Kit (see Supplementary Materials S5).

#### 2.3.2. Incubation of carbon-enriched marine-derived microbial communities with plastic-derived leachates

A schematic depicting our experimental set-up is provided in Figure S3. Carbon-enriched marine-derived microbial communities were exposed to the four plastic leachates and an ASWS control at two concentrations: (1) 35 ml each of leachate and culture (designated the ’50% carbon’ group) and (2) 17.5 ml of leachate, 17.5 ml fresh Marine Broth 2216 and 35 ml culture (designated the ’75% carbon’ group). Both treatments included four replicates for each plastic type (LLDPE, PA6, PET, PLA) and an uninoculated control, except for PA6 and PET 75% carbon treatments with only three experimental replicates due to limited available glassware. Both 50% and 75% carbon treatments were incubated for seven days at room temperature (18 – 22°C) in Erlenmeyer flasks in a rotary incubator.

After incubation, 20 ml aliquots were collected for RNA and DNA analysis. Cell pellet samples for RNA analysis were snap-frozen using liquid nitrogen and stored at −80°C. Those for DNA analysis were stored at −20°C. The remaining inoculated solutions were pelleted via centrifugation (3756 x g for 10 minutes) to be used as inoculum for a 0% media carbon group, while the supernatant was snap-frozen in liquid nitrogen and stored at −80°C for GC-MS analysis.

#### 2.3.3. Adding plastic leachate to marine-derived microbial communities with no additional media carbon

Pelleted cells for the ‘0% carbon’ group were washed in 5 ml of 1 x PBS, pH 7.4, and centrifuged at ∼3756 x g for 10 minutes, three times, to remove residual media. Cells were then used as inoculum for incubation with plastic leachate-only media (with no additional carbon source). Washed cells, previously grown in 50% carbon, were used to inoculate four replicates of each plastic leachate type (LLDPE, PA6, PET, PLA), along with an ASWS control. Each plastic type also included an uninoculated control sample. The inoculated plastic leachate was shaken at room temperature (18 - 22°C) for seven days, along with uninoculated controls. Material for DNA, RNA and GC-MS analysis was collected as previously described.

### 2.4. Gas chromatography-mass spectrometry (GC-MS) analysis of organic compounds in the plastic leachates

The organic chemical composition of each plastic leachate type was assessed using a non-targeted approach by solvent extraction gas chromatography-mass spectrometry (GC-MS) as detailed in the Supplementary Materials (S6).

### 2.5. DNA extraction, metabarcoding and amplicon sequencing

Microbial DNA was extracted from the centrifuged pellets of each sample using a DNeasy PowerSoil Pro kit (Qiagen, Hilden, Germany). Polymerase chain reactions (PCRs) were performed to amplify (i) the hypervariable V3/V4 region of the bacterial small subunit of the ribosomal RNA (16S rRNA) gene using the universal amplicon primer pair 341F and 785R and (ii) the fungal internal transcribed spacer 2 (ITS2) region of the nuclear ribosomal gene using the primer pair fITS7 and ITS4 (Table S2). Fungal primer reactions gave low yields, and replicate PCR products were subsequently pooled before sequencing. DNA sequencing was completed on an Illumina MiSeq instrument. Additional methods for DNA extraction, amplification and sequencing are provided in Supplementary Materials (S7).

### 2.6. RNA extraction and sequencing

Since the community DNA analysis showed the greatest difference in microbial composition following PLA leachate exposure, the transcript expression of PLA leachate-exposed communities was compared with the preconditioned and ASWS-supplemented communities. Microbial RNA was extracted from 20 ml pelleted samples using an RNeasy PowerBiofilm kit (Qiagen, Hilden, Germany). The Auckland Genomics Facility conducted RNA sequencing by ligation with Ribo-Zero Plus, using an Illumina Stranded Total RNA Prep kit (Cat. No. 20072063), and sequencing on an Illumina NovaSeq X instrument. Additional methods for RNA extraction and sequencing are provided in the Supplementary Materials (S8).

### 2.7. Accession Numbers

Raw sequence reads are available from the NCBI Sequence Read Archive (SRA) under the submission number SUB14024939.

### 2.8. Processing and analysis of DNA sequence data

Sequence adapters from reads were removed with Trimmomatic (version 0.39) (Bolger et al., 2014), using the parameters: ILLUMINACLIP: 2:40:15. Primers were removed using Cutadapt (Martin, 2011). Successfully trimmed reads were processed using the DADA2 pipeline (Callahan et al., 2016) in R (version 4.0.1) (R Core Team, 2020) following package instructions. Samples were dereplicated and pooled together for amplicon sequence variant (ASV) inference. Paired-end reads were merged, chimeric sequences removed, and an ASV table was generated. Taxonomic assignment was performed against the SILVA reference database for bacterial 16S rRNA sequence reads (version 138) (McLaren, 2020) and against the UNITE reference database for fungal ITS sequence reads (general FASTA release version 10.05.21) (Abarenkov et al., 2020) using the native implementation of the naïve Bayesian classifier method. ASVs were defined as DNA sequences sharing 100% identity.

### 2.9 DNA amplicon bioinformatics and quantitative analysis

All quantitative and statistical analyses and data visualisation were performed using R (version 4.2.2) (R Core Team, 2020) as described in the Supplementary Materials (S9), using PERMANOVA and non-metric multidimensional scaling (nMDS) ordinations based on microbial community Bray-Curtis dissimilarities (rarefied ASVs were randomly subsampled to the minimum non-chimeric library size - rarefying; set.seed(123), bacteria n = 1151, fungi n = 1469).

To determine the presence of any previously reported putative plastic degraders in our plastic leachate samples, bacterial and fungal ASVs were BLAST-searched (Camacho et al., 2009) against the PlasticDB database of putative plastic-degrading microorganisms (www.plasticDB.org) (Gambarini et al., 2022). Potential degraders were identified at the genus and species levels.

### 2.10. Meta-transcriptome assembly and analysis

Transcriptomic sequences were processed as described in the Supplementary Material (S10). After quality checking, ribosomal RNA was removed from the dataset, the remaining reads *de-novo* co-assembled and clustered based on 95% confidence matching isoforms before the metatranscriptome was annotated. Finally, transcripts were searched using DIAMOND blastp against a plastic-degrading protein database, PlasticDB (Gambarini et al., 2022). We used multiple packages to assess differential expression among sample groups, as also described in the Supplementary Material (S10).

## 3. Results

A total of 68 samples were processed for bacterial 16S rRNA and fungal ITS gene sequence analysis (five leachate types [PLA, LLDPE, PA, PET, ASWS] x three carbon conditions [0%, 50%, 75%] x four replicates + five pooled media samples + three pre-conditioned samples = 68). Experimental controls had low sequence read counts and were subsequently removed from downstream analysis. After quality filtering, 2075 and 100 unique ASVs were obtained for bacteria and fungi, respectively. Following rarefaction, 912 bacterial and 74 fungal unique ASVs were retained for further analysis.

### 3.1. Organic component analyses by GC-MS

No distinct GC-MS data clustering based on Bray-Curtis dissimilarity was observed based on leachate type alone, although PLA 0% carbon samples clustered independently from all other samples (Figure S4). Toluene was prevalent in all four plastic leachates (Figure S5), but most organic components were only found at relatively low concentrations (e.g. Pyrrolo(1, 2-a)pyrazine-1, 4-dione, hexahydro). L-lactic acid, 1,2-benzenedicarboxylic acid, and 3-nitro were identified in PLA experimental replicates as well as controls where there was no bacterial community present, indicating abiotic PLA degradation (Figure S5D). Caprolactam was identified in PA experimental replicates and controls, with cyclopentane-carboxylic acid and 1,8 diazacyclotetradecane −2, 9-dione found in the 0% PA carbon media group (Figure S5B). LLDPE and PET had the lowest concentrations of organic substances in the leachate of any plastic leachate type. LLDPE leachate contained pyrrolo[1,2-a]pyrazine-1,4-dione, hexahydro (97% best hit), hexadecanoic acid, methyl ester (98% best hit) and 9-octadecenoic acid (Z), methyl ester (99% best hit), with no discernable distinction between media carbon percentages (Figure S5A). PET leachates contained two organic components, 2(5H)-furanone, 3,5,5-trimethyl (64% best hit) or 2-(1-hydroxyethyl)-2-cyclopenten-1-one (64-59% best hit), and Z-3-methyl-2-hexenoic acid (43% best hit) or methyl (2E)-4-oxo-2-pentanoate (38% best hit) (Figure S5C), as the most abundant organic components specific to PET. Salicylic acid, 2TMS derivatives were only in ethyl acetate controls (Figure S5A). No organic components related to plastic degradation were specifically dominant in LLDPE or PA leachate samples (*P* > 0.05).

### 3.2. Cell Viability

Permutational ANOVA revealed that cell viability was significantly influenced by the percentage of media carbon and duration of the experiment (*P* = 0.002, *P* = 0.001, respectively) but not plastic leachate type (*P* = 0.385) (Figure S6). The proportion of viable cells decreased from day one to day nine for LLDPE, PET and ASWS of the 50% carbon group, indicative of cell death (Figure S6). In contrast, for the same carbon group, PLA and PA show an increase, which is likely caused by the use of carbon substrates within the plastic leachate (Figure S6 & S7). Cell viability in the 0% carbon treatments, observed at seven days only, was significantly lower than that of the 50% and 75% carbon treatments (permutational ANOVA *P* > 0.01); similar trends were observed for the OD_600nm_ data (Table S3).

### 3.3. Effects of leachate type on bacterial community composition

All data from bacterial communities exposed to PLA-based leachates clustered independently from other community data (PERMANOVA *P* = 0.001), indicating that they were distinct from LLDPE, PA and PET leachate-exposed communities (Figure 1A). This is supported by pairwise PERMANOVA (when compared to PLA: ASWS *P* = 0.001, LLDPE *P* = 0.001, PA *P* = 0.001, PET *P* = 0.001, preconditioned (Time 0) *P* = 0.007). Analysis of the phyla contributing most to community differences revealed that the higher relative abundance of *Proteobacteria* in PLA leachate-exposed samples was the largest factor influencing the observed difference (Figure 1C). While there was a high degree of overlap among data from the other plastic leachate types, pairwise PERMANOVA revealed that PA-exposed samples were distinct from LLDPE-exposed and pre-conditioned samples (*P* = 0.05 and *P* = 0.03, respectively). Weakly significant differences were also detected when comparing PA-exposed samples to PET-exposed or ASWS samples (*P* = 0.08 and *P* = 0.07, respectively). No other pairwise plastic-type comparisons were significant. However, PERMANOVA revealed significant differences between carbon condition types (*P* = 0.001) (Figure 1B). The 0% carbon communities were distinct from all other carbon concentrations (Pairwise *P* = 0.001), and 50% was distinct from 75% (*P* = 0.002).

**Figure 1.**
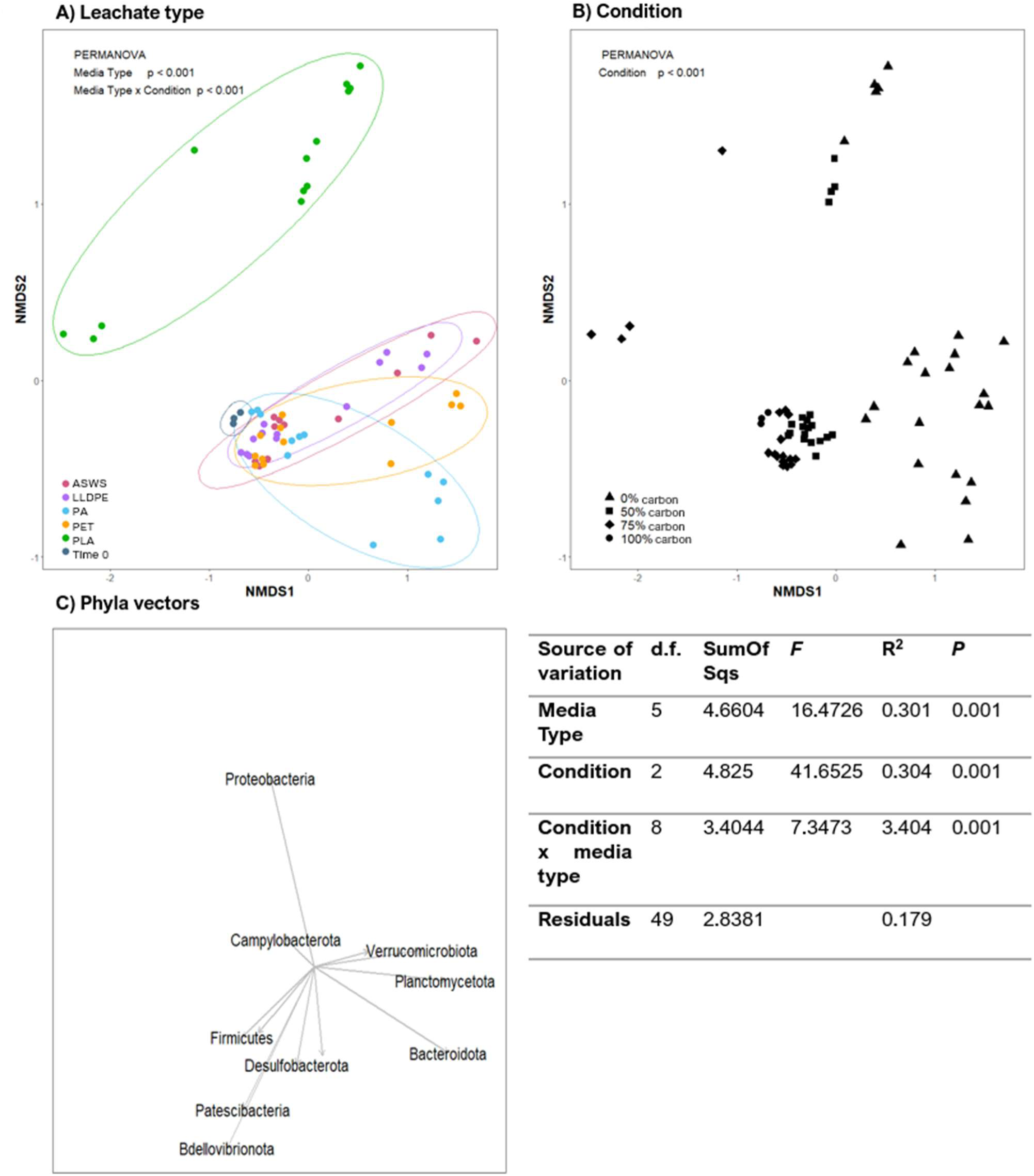
Changes to bacterial community composition following exposure to plastic leachates. Non-metric multidimensional scaling (nMDS) ordination of Bray-Curtis similarities for (A) bacterial community data grouped by media type ((LLPDE) linear low-density polyethylene, (PA) polyamide-6, (PET) polyethylene terephthalate, (PLA) polylactic acid, (ASWS) artificial sea water salts broth, (Time 0) a control Marine Broth 2216 medium aged under the same conditions before leachate addition. (B) Effect of plastic leachates stratified by growth media carbon condition. Ellipses were drawn using the geom_mark_ellipse function in the R package ggforce, assuming a multivariate t-distribution. (C) The dominant phyla driving the compositional differences in plots A & B are denoted by the arrow length in plot C, representing significance at *p* < 0.05, in scale with A & B, as determined by the linear regression model of the vegan package envfit function in R.The results of a PERMANOVA quantifying variation in the data as attributable to either media type, condition and their interaction are presented in the bottom right quadrant of this figure.

### 3.4. Taxonomic variation in bacterial and fungal community composition across leachate type and carbon concentration

As with cell viability, the relative abundance of bacterial taxa was significantly impacted by carbon condition (PERMANOVA, *P* = 0.001). Relatively higher abundances of *Bacteroidota*, *Planctomycetota* and *Verrucomicrobiota* phyla were most associated with the 0% carbon condition (Figure 1), despite *Proteobacteria* being the most dominant phyla across all conditions (Figure S10). Weak differences were detected when comparing either 50% and 75% treatment communities to the preconditioned community (100%) (*P* = 0.06 and *P* = 0.08, respectively), although *Bdellovibrionota* and *Patescibacteria* were more abundant under the 75% carbon conditions. However, relative abundance was also influenced by leachate media type. In particular, *Bacteriodota* was relatively more abundant in the PA-exposed 0% carbon media-type samples, and PLA-exposed leachate communities remained distinct from other leachate-exposed communities. PLA-exposed leachate communities were dominated by *Proteobacteria,* with *Bacteroidota* only present at relatively low abundances (Figure 1C, Figure S10).

At the family level, LLDPE, PA, and PET-exposed communities shared similar taxa relative abundances, while PLA-exposed communities remained distinctive (Figure 2). PLA 0% and 50% carbon were relatively similar in community composition, both dominated by the proteobacterium, *Halomonadaceae*; all other plastic leachates and control samples contained <1% *Halomonadaceae*. Additionally, unlike other leachate-exposed samples, PLA 0% and 50% carbon communities had a high abundance of *Aeromonadaceae*. PLA 0% carbon also contained between 4% and 13% *Stappiaceae*, found in all other samples at <1% abundance. PLA 75% was the only sample to be dominated by *Vibrionaceae*. *Thalassospiraceae*, *Sneathiellaceae*, *Flavobacteriaceae* and *Rhodobacteraceae* were more abundant in ASWS, LLDPE, PA, and PET 0% carbon samples than other carbon conditions. PET leachate-exposed communities were highly abundant with *Thalassospiraceae,* and PA abundant with *Rhodobacteraceae* and *Thalassospiraceae*. Most ASVs were found at relatively low abundance across various samples and replicates. Overall, there was a shared change in taxonomically classified ASVs when comparing the 0% carbon treatment with other carbon treatments, except for PLA. Similar findings could be observed at the phylum level (see Figure S10) and for the fungal data, where the greatest differences were observed for the PLA-leachate-exposed communities (Supplementary Materials S11 and Figure S11).

**Figure 2.**
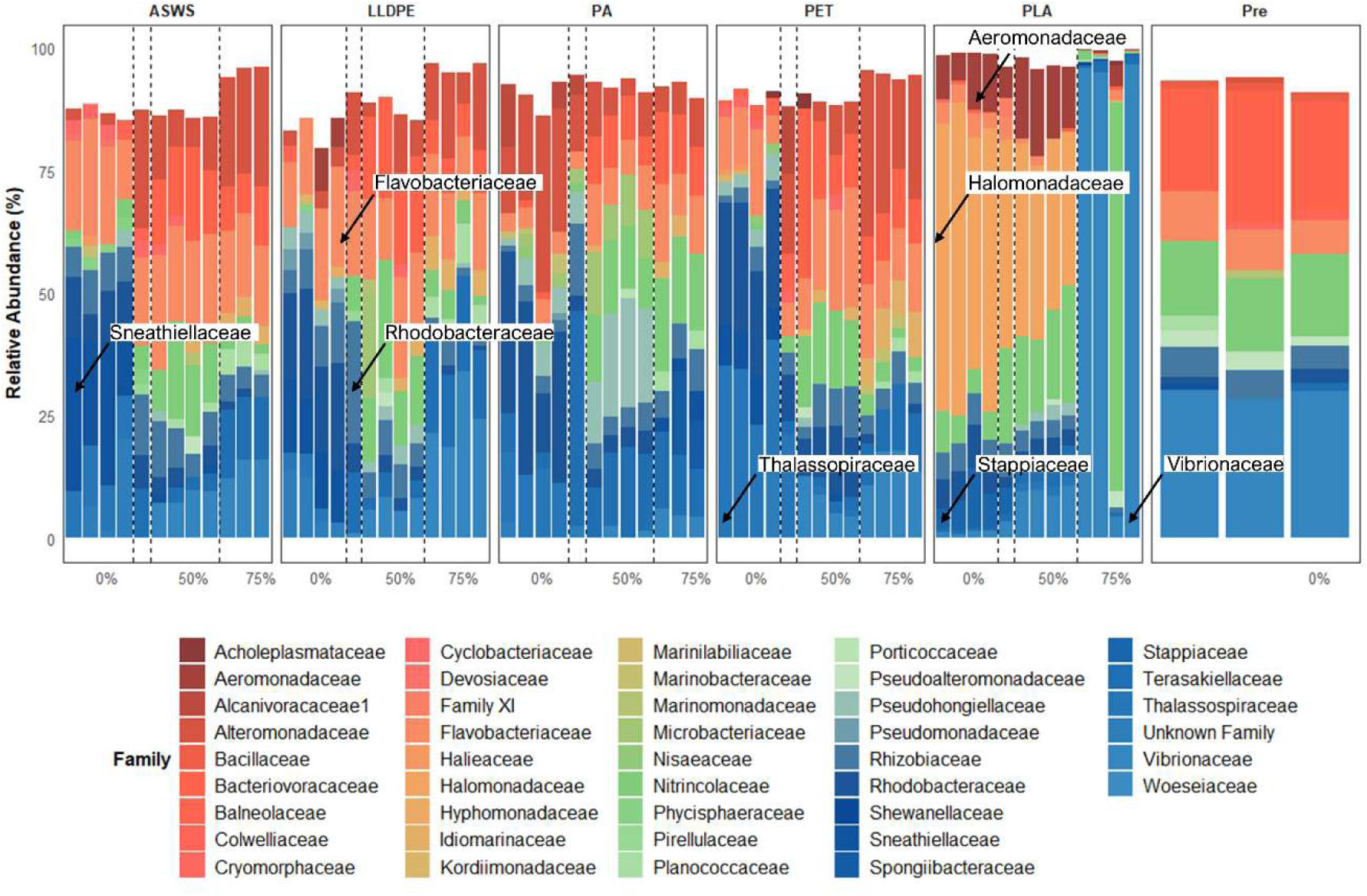
Bacterial relative abundance plots, based on analysis of 16S rRNA genes of the top 10 families from each rarefied plastic leachate sample. Samples include (ASWS) artificial seawater broth, (LLDPE) linear low-density polyethylene, (PA) polyamide-6, (PET) polyethylene terephthalate, (PLA) polylactic acid, and (Pre) precondition growth media [Marine Broth 2216]. Following filtering and rarefaction, sequence reads were identified using the DADA2 package in R (R Core Team, 2020) for read processing, and a non-redundant SILVA_v138.1 database (McLaren, 2020) for taxonomic assignment for samples which contained at least 1000 sequence reads. The dotted lines indicate treatment groupings, with a 0% replicate made from the 75% carbon group as inocula. Data relating to one ASW-75, one ASW-0 and one PA-75 incubation were removed due to the low read counts.

### 3.5. Screening for reported putative degraders

A total of 17 bacterial genera identified in this study were related to genera with reported plastic-degrading phenotypes (Figure 3). The putative plastic biodegraders identified in this study have been reported to degrade a range of plastic substrates, indicating wide substrate specificity. For instance, *Alcanivorax* spp. (Khandare et al., 2021; Sekiguchi et al., 2011), *Bacillus* spp. (Adıgüzel and Tunçer, 2017; Kumari et al., 2019; Sekiguchi et al., 2011), *Pseudomonas* spp. (Deepika and Jaya, 2015; Sekiguchi et al., 2011) and *Vibrio* spp. (Danso et al., 2018) have all been reported to degrade LDPE and PCL, among other plastics. Most putative plastic degraders were found in at least one sample from each leachate treatment. However, *Halomonas* spp., reported to degrade LDPE (Khandare et al., 2021), was found to be the most abundant of any previously reported plastic degraders, dominating PLA 0% and 50% carbon samples, making up as much as 61% of the ASVs identified (Figure S12). While at low abundances, other leachate-exposed 0% carbon samples had distinctive reported degraders. *Leucobacter* sp. was found at about 5% relative abundance in PA 0%, in addition to PA 50% carbon samples. PA 0% also had *Alcanivorax* spp. at unusually high abundance, as much as 10% of the relative taxa abundances in select samples (Figure S12). Only five bacterial ASVs reported as putative plastic biodegraders were identified at the species level: *Lysinibacillus sphaericus*, *Lysinibacillus xylanilyticus*, *Nesiotobacter exalbescens*, *Pseudomonas pachastrellae*, and *Vibrio parahaemolyticus*. All species were found at < 3% relative abundance across all leachate-exposed samples, with *L. xylanilyticus* most abundant at 2.7% in LLDPE 75% carbon samples.

**Figure 3.**
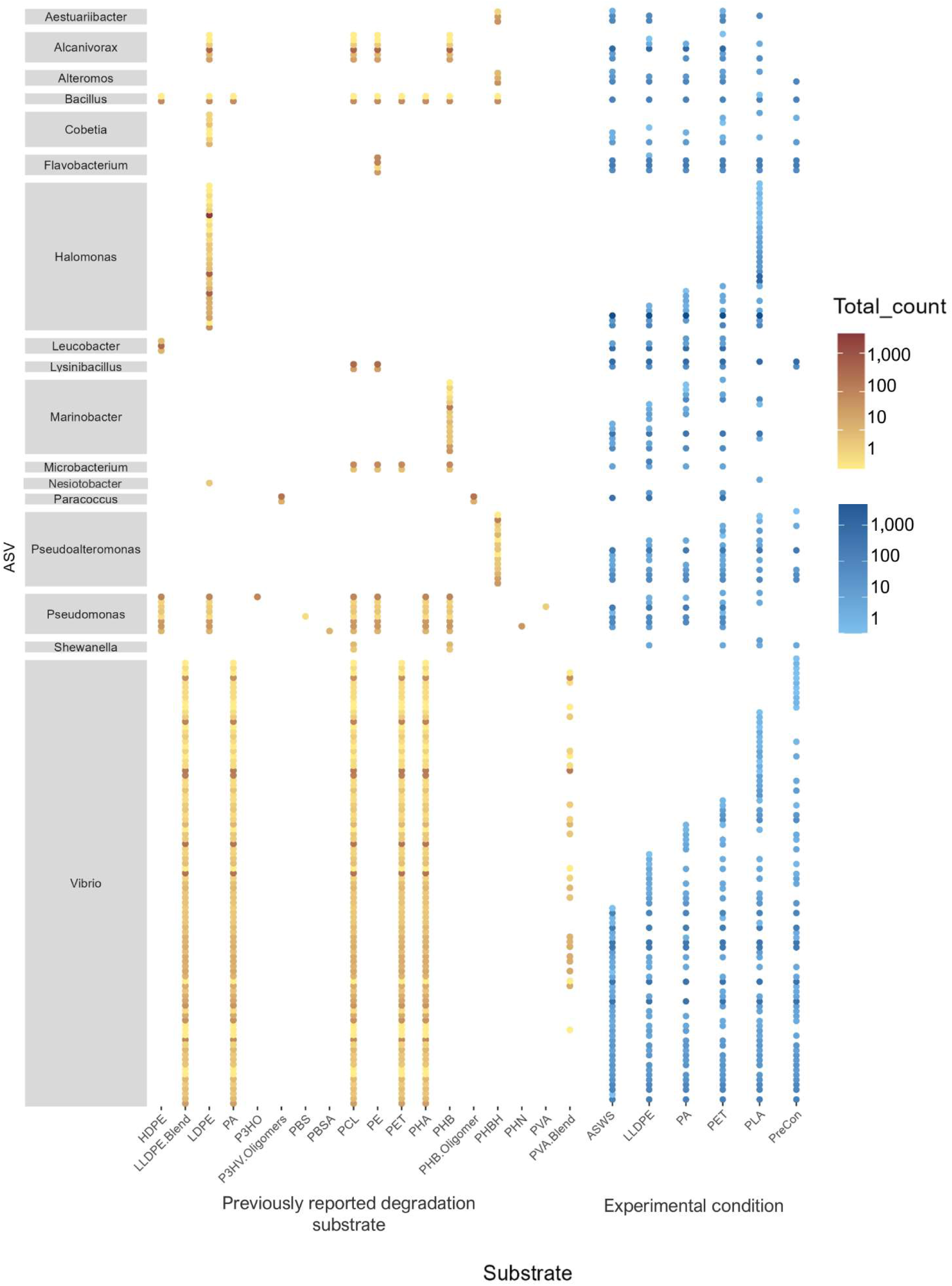
Log scaled total ASV count assigned to genera of previously reported as putative plastic degraders, identified using PlasticDB (accessed Nov 2022) (Gambarini et al., 2022). The plot on the left (orange/yellow) illustrates the total number of ASVs in the dataset previously reported to degrade different plastics grouped by genera. The plot on the right (blue) contains the same ASV data, quantified by their abundance in different experimental treatments in which the ASVs were found.

Across the 100 unique fungal ASVs, 11 were identified as belonging to genera previously reported as putative plastic biodegraders before sample filtering and rarefaction (Table S4). However, only five of these ASVs were retained post-filtering and rarefaction: *Coniochaeta* sp., *Cryptococcus* sp., *Fomitopsis* sp., *Fusarium* sp., and *Rhodotorula* sp.. *Coniochaeta* sp., *Fomitopsis* sp., and *Fusarium* sp. have been reported to degrade PU (Navarro et al., 2021), with *Fusarium* sp. also reported to degrade LDPE (Das and Dash, 2014), PHB (Jeszeová et al., 2018), and nylon (Tachibana et al., 2010). *Cryptococcus* sp. has been reported to degrade PBS, PCL, PHB, PLA, and PHA (Masaki et al., 2005), and *Rhodotorula* sp. reported to degrade PLA (Nair et al., 2016). All fungal ASVs identified as previously reported putative plastic degraders were <1% of the relative abundance within each sample, except for *Coniochaeta* sp., which was 22.3% in PET-75.

### 3.6. Plastic leachate-specific bacterial indicator ASVs

Indicator value analyses (IndVal) revealed that certain bacterial ASVs were significantly associated with different plastic leachate media types (Figure S13). A total of 104 out of the 912 unique ASVs are potential indicators, by both specificity and fidelity, of each media type. Most indicator ASVs (n = 27) were assigned to PLA, 8 to ASWS, 12 to PA, two to PET and only one to LLDPE. LLDPE, PET and PLA only contained indicators from the phylum *Proteobacteria*. Most ASVs in PLA were assigned to Proteobacteria (mostly *Halomonas*). No indicator ASVs were assigned to a plastic biodegrader reported to degrade plastic for the same leachate type.

### 3.7. Transcriptional activity of taxa exposed to PLA leachate focusing on genes encoding putative plastic-degrading enzymes

Analysis of the meta-transcriptome sequences, as compared to sequences within the PlasticDB database (Gambarini et al., 2022), revealed that among the transcripts related to plastic-degradation, those encoding for PLA degradation were very common (Figure S14). However, only nine transcripts matched the database with above a 90 % identity match, three of which were identified as esterases reported to degrade PLA. The remaining six transcripts were identified as PHB depolymerase, polyester hydrolase, 3HV dehydrogenase and esterases described to degrade polyurethane and polycaprolactone.

A total of 650,500 transcripts were differentially expressed in PLA samples compared to ASWS-exposed and preconditioned samples (Figure S15). Of the transcripts that were found in at least two of the models with an FDR adjusted P-value < 0.05 and a positive log fold change, indicating a proportional increase in PLA samples, three transcripts were found in PLA 50% media carbon samples, 18 in PLA 75% media carbon, and 74 in PLA 0% media carbon (Table S5). None of the sequenced transcripts were identified as coming from specific PLA degrading genes. Most transcripts had annotated functions associated with PEG aldehyde dehydrogenase, found in all conditions, and greater abundance in PLA-leachate-exposed samples. PETase-assigned transcripts were also prevalent in PLA 0% media carbon samples.

Since our amplicon data revealed *Halomonas* sp. to be prevalent in PLA-leachate-exposed communities, we aligned our transcript reads to 13 *Halomonas* spp. reference genomes available on NCBI at the time of this study (12/12/2024). Two of the reference genomes contained comparatively strong alignments for PLA-exposed community transcripts. Up to 49.69 % and 32.04 % of total transcript reads aligned with *H. hydrothermalis* Slthf2 (Takeyama et al., 2020) and *H. qaidamensis* XH36 (Zhang et al., 2022) reference genomes, representing 14.87 M and 7.74 M transcript reads respectively. Neither reference genome contained known PLA-degrading genes catalogued within PlasticDB. Instead, both genomes contained several hits at <50% identity match for other plastic degrading genes including esterases, hydrolases, laccases, polyesterases and PHB depolymerases. However, of the 36 differentially expressed transcripts, when comparing PLA-exposed communities and ASWS control communities, no transcripts were identified as being related to genes previously reported to code for putative plastic degradation.

## 4. Discussion

In this study, we identify changes in marine microbial communities after exposure to the leachates of relatively inert and additive-limited plastics. Understanding the impact these plastic leachates have on marine or other microbial communities has important implications for our understanding of the lingering effects of plastic waste on ecosystem functioning.

### 4.1. Plastic leachate composition

Both PET and LLDPE require exceptionally high temperatures to thermally degrade (150-300°C) or the presence of catalysts involved in the breakdown of ester or C-H bonds (Lear et al., 2022). In this study, both plastics were found to leach very few substances into the surrounding media. Indeed, in a dynamic leaching experiment investigating temperature and time, Bridson et al., (2023) found that LLDPE and PET released cumulative additives up to five orders of magnitude less than PA6. We found a similar trend, with GC-MS detection peaks for LLDPE and PET essentially indistinguishable from the ASWS-exposed community. However, the PA6 leachate contained the oligomer caprolactam, indicating the presence of PA6 monomers. The monomer is likely a residual artefact of incomplete polymerisation (Klaeger et al., 2019), still found in the polymer formation but not bound to the structure itself. Therefore, to convincingly identify polymer degradation, additional chemical analyses would be necessary to confirm the absence of the monomer prior to leaching. These three plastics (LLDPE, PET and PA6) were distinct from PLA, which has a much lower glass transition temperature, closer to 55°C (Nguyen et al., 2018). With a low glass transition temperature, we expected a higher proportion of associated monomers (L-lactic acid) and other additive chemicals in the PLA-leachate, as complete PLA conversion occurs during hydrolysis (Cristina et al., 2018). As expected, we found L-lactic acid in all PLA leachate samples. The release of these oligomers/monomers from widely used plastics such as PLA and PA6 has significant implications for impacting microbial community functioning.

### 4.2. Microbial community patterns in leachate-exposed communities

Microbial growth and community composition were significantly influenced by both the plastic leachate type and the media carbon concentration, with the strongest differences occurring in PLA- and PA6-exposed communities with no additional media carbon. Previous studies examining plastic-derived leachates have shown that leachates cause specific changes in microbial ecosystem functioning, such as those related to antibiotic resistance and virulence (Vlaanderen et al., 2023), and photosynthetic ability (Sarker et al., 2020; Tetu et al., 2019). In polyethylene-based plastic leaching experiments, Romera-Castillo et al., (2018) demonstrated that different polyethylene-based plastic types did not negatively impact bacterial growth rates. We observed similar results. Viable cell counts of LLDPE and PET-exposed-leachate communities were essentially indistinguishable from control samples without leachate, showing similar live cell counts to controls during the microbial incubation period, even observing viable cells in 0% media carbon after seven days. PA6 and PLA showed different trends, increasing viable cell counts even in 0% media carbon. This is likely due to oligomers, L-lactic acid and caprolactam leaching from PLA and PA6, respectively, supporting microbial growth. Caprolactam is known to inhibit bacterial growth in several genera, namely *Bacillus* sp. and *Rhizobium* sp., while promoting the growth of caprolactam-degrading bacteria (Baxi, 2013), and even improves oxygen production in *Desmodesmus quadricauda* algae (Kalinová et al., 2016). Similarly, L-lactic acid can both inhibit and promote growth in select bacteria. Lactic acid has been reported to inhibit the growth of pathogenic taxa such as *Salmonella* Enteritidis, *Escherichia coli*, and *Listeria monocytogenes* (Wang et al., 2015) by inhibiting the synthesis of, or even destroying cellular proteins, resulting in cell death (Zeng et al., 2010). However, several taxa can metabolise and ferment lactic acid (Holzapfel and Wood, 2014). Most other chemicals detected in the leachates, such as toluene, have also been shown to be degraded by various microorganisms (Parales et al., 2008), potentially supporting microbial growth, in addition to the oligomers found in PLA and PA6. This suggests that these two oligomers, in conjunction with other chemicals found in PLA and PA6, have the potential to shape the community composition, which was observed in our study to varying degrees.

PLA- and PA6-exposed leachate community composition differed from those exposed to LLDPE and PET. In all conditions, PLA leachate-exposed bacterial communities were almost entirely dominated by Proteobacteria, specifically the family Halomonadaceae. This bacterial group, commonly found in marine environments, has promising biotechnological applications, mainly due to its high salt tolerance, nonspecific growth requirement (de la Haba et al., 2011) and demonstrable roles in bioremediation (Piubeli et al., 2012). While it is suspected to have a functional role in LDPE degradation (Khandare et al., 2021), the PLA leachate may generate conditions such as low pH that exclude the growth of other bacteria while promoting *Halomonadaceae* growth. However, differential transcriptome analysis could not identify areas of the genome that could explain its dominance within the community (e.g., relating to neutralising or exporting excess cellular protons). PLA leachate-exposed fungal communities were distinct from other leachate-exposed communities, containing a range of taxonomic families not observed in other leachate-exposed communities. This could result from the more complex chemical composition of PLA leachate, providing more nutrients for growth. However, with the lack of fungal sample replicates, we cannot confirm these trends. Similarly, PA6-exposed bacterial communities contained several distinct taxa, namely Actinobacteriota, specifically *Microbacteriaceae*, which was not observed in other communities. While the exact function of *Microbacteriaceae* identified in our study may be unknown due to the wide variety of genera that exists (Evtushenko and Takeuchi, 2006), the family does contain genera previously reported to degrade plastic, namely *Leucobacter* spp. (Devi et al., 2019) and *Microbacterium* spp. (Suzuki et al., 2017) which were both found in PA6-exposed communities and reported to degrade HDPE (Devi et al., 2019) and PET (Yan et al., 2021), respectively. Unlike PLA-exposed communities, there were few differences between communities exposed to PA6 leachate and leachate-free controls.

### 4.3. Previously reported putative plastic degraders and indicator species

We detected relatives of putative plastic biodegraders within the different plastic leachates at low environmental abundances in nearly all leachate conditions. Species belonging to the genera *Pseudomonas* (Molitor et al., 2020; Montazer et al., 2018), *Bacillus* (Aravinthan et al., 2016; Giacomucci et al., 2019; Khandare et al., 2021), and *Vibrio* (de Vogel et al., 2021; Sarkhel et al., 2020), have been reported to degrade or have the potential to degrade multiple types of synthetic polymers. Specifically, an ASV assigned to *Pseudomonas pachastrellae,* found in a range of leachate conditions, has been reported to degrade PS (Kim et al., 2021), LDPE (Nadeem et al., 2021), PET (Bollinger et al., 2020), PU, and PCL (Molitor et al., 2020). At the same time, an ASV assigned to *Vibrio parahaemolyticus* is reported to degrade PCL, PET (Danso et al., 2018), nylon (Sudhakar et al., 2008) and LLDPE blends (Raghul et al., 2014). However, these genera are ubiquitous and typically found in marine environments; therefore, they are likely abundant in any sample without actively degrading plastic-related substances. Additionally, we found *Lysinibacillus xylanilyicus* and *L. sphaericus* in all samples; they are reported to degrade PE (Sangale et al., 2019; Shahnawaz et al., 2016a, b) and PCL (Xu et al., 2007). Again, given their ubiquitous presence in all experimental samples, we would not expect these species to necessarily be involved in each of these plastic degradation-associated pathways.

Several PLA-, PA- and ASWS-exposed communities contained some indicator ASVs related to putative plastic biodegraders. These included *Flavobacterium* spp. in ASWS-exposed communities, *Alcanivorax* spp. and *Leucobacter* spp. found in PA-exposed communities, and *Halomonas* spp. and *Vibrio* spp. found in PLA-exposed communities. However, these genera, except *Vibrio* spp. and *Alcanivorax* spp., have previously been reported to degrade polyethylene-based plastics (Devi et al., 2019; Khandare et al., 2021; Koutny et al., 2009) and appear at low abundances. It is unclear if these taxa are involved in plastic leachate degradation or carbon assimilation. Still, indicator taxa in PA6 leachate communities could have some pathway function involved in PA6 leachate degradation or carbon assimilation. Conversely, *Halomonas* spp. were unusually abundant in PLA leachate-exposed communities, indicating some relationship with PLA-leached chemicals. *Halomonas* strains have previously been reported to degrade aromatic hydrocarbons (Delacuvellerie et al., 2019; Govarthanan et al., 2020), which are typically used in the synthesis of plastics, but there are no reports of *Halomonas* strains degrading PLA or PLA-derived chemicals. Toluene, a common aromatic hydrocarbon, was found in all plastic leachate samples, yet the same abundances of *Halomonas* spp. were not noted in all plastic leachate exposed communities. This indicates that *Halomonas* spp. may not have been involved in toluene degradation but may have a role in the biodegradation of other PLA leachate-associated chemicals or that PLA leachates create an environment that enriches for *Halomonas* spp.. However, differential expression analysis of the meta-transcriptome did not reveal any transcripts to explain *Halomonas’s* role within the community.

Like bacteria, fungi related to putative plastic biodegraders were not necessarily involved in plastic leachate biodegradation. Most ASVs were found to be too low in abundance to significantly impact the community functioning. Despite this, *Bjerkandera adusta*, found in PA6 and PLA leachate-exposed communities, was previously reported to biodegrade both PLA (Jeszeová et al., 2018) and nylon-6 (Friedrich et al., 2007). Additionally, *Fusarium keratoplasticum*, found in PA6 leachate-exposed communities, was reported as a nylon-4 degrader (Tachibana et al., 2010). Both fungal species are commonly found in soil or plant microbiomes, where complex natural polymers are found at higher densities than in marine environments. We might expect these species to be active in polymer degradation in soil environments.

### 4.4. Differential expression of transcripts in PLA-leachate-exposed communities

Understanding metabolic activity is arguably the most important facet of understanding how microbial communities change due to different stressors. While many differentially expressed transcripts were found in the PLA leachate-exposed communities, a relatively low number of these transcripts were associated with previously reported plastic biodegradation proteins. This is likely the result of (i) plastic-degrading proteins still being relatively poorly understood and (ii) that proteins being expressed and regulated in these communities may not be directly linked to polymer biodegradation but may have roles associated with monomer degradation or other chemicals associated with plastics leachates, or (iii) that some plastic-degradation encoding genes may be constitutively expressed. Of the transcripts mapped to genes encoding plastic-degrading proteins, there was a surprising lack of specific PLA-degrading proteins. Instead, the transcripts were dominated by those encoding esterases and dehydrogenases. Esterases are widely described in plastic degradation, with novel esterases identified through metagenomic sequencing that are predicted to hydrolyse PLA (Hajighasemi et al., 2018). Conversely, plastic-degrading dehydrogenases are less widely cited, and their role is often associated with a specific substrate, e.g., polyethylene glycol (PEG) and PHA biodegradation (Kasuya et al., 1999; Ohta et al., 2005). Therefore, testing of enzymatic activity on specific leachate components is required to fully understand these protein functions in plastic-associated communities. A deeper analysis of the degradation pathways for all plastic-associated chemicals is required to fully understand how the communities respond to different plastic leachates.

### 4.5. Conclusion

With marine environments exposed to different types of plastic waste, it is essential to understand how plastic deteriorates over time and how plastic leachates affect ecosystem functioning. Waste plastics exposed to harsh abiotic factors will likely fragment and leach functional and non-intentionally added additives into the environment, which could impact the marine microbiome. The research presented here shows that while inert unadulterated plastics are unlikely to release non-intentionally added chemicals into the marine environment via thermal and hydrolytic deterioration, biodegradable plastics and, more interestingly, polyamides are likely to deteriorate over time, releasing a complex consortium of chemicals. These chemicals can alter the surrounding microbiota, selecting microbes that can tolerate or utilise these chemicals. Further investigation into the exact functions of these microbes is still required. The use and management of even biodegradable plastics and polyamides must be treated with caution to avoid disturbing the delicate equilibrium of the marine ecosystem.

## Acknowledgements

He mihi ki ēnei hinonga pakihi, kaunihera rātou ko ngā tangata whenua mō ā rātou awhina mai. We thank the Viaduct Harbour for allowing the deployment of, and access to, the study structure to collect environmental samples. We acknowledge Ngāi Whātua Ōrākei who express cultural authority over Tāmaki Makaurau-Auckland.

## Funding

This study was supported by the New Zealand Ministry of Business, Innovation and Employment, Endeavour Research Programme fund C03X1802 (Impacts of microplastics on New Zealand’s bioheritage systems, environment, and eco-services).

## Notes

### Competing Interest Statement

The authors have declared no competing interest.

## References

Abarenkov, K., Zirk, A., Piirmann, T., Pöhönen, R., Ivanov, F., Nilsson, R., Kõljalg, U., 2020. UNITE general FASTA release for Fungi UNITE Community 761.

Adıgüzel, A.O., Tunçer, M., 2017. Purification and characterization of cutinase from Bacillus sp. KY0701 isolated from plastic wastes. Preparative Biochemistry Biotechnology Advances 47, 925–933.

Aravinthan, A., Arkatkar, A., Juwarkar, A.A., Doble, M., Biotechnology, 2016. Synergistic growth of *Bacillus* and *Pseudomonas* and its degradation potential on pretreated polypropylene. Preparative Biochemistry 46, 109–115.

Baxi, N.N., 2013. Influence of ε-caprolactam on growth and physiology of environmental bacteria. Annals of Microbiology 63, 1471–1476.

Bolger, A.M., Lohse, M., Usadel, B., 2014. Trimmomatic: a flexible trimmer for Illumina sequence data. Bioinformatics 30, 2114–2120.

Bollinger, A., Thies, S., Knieps-Grünhagen, E., Gertzen, C., Kobus, S., Höppner, A., Ferrer, M., Gohlke, H., Smits, S.H., Jaeger, K.-E., 2020. A novel polyester hydrolase from the marine bacterium *Pseudomonas aestusnigri*–structural and functional insights. Frontiers in Microbiology 11, 114.

Bridson, J.H., Abbel, R., Smith, D.A., Northcott, G.L., Gaw, S., 2023. Release of additives and non-intentionally added substances from microplastics under environmentally relevant conditions. Environmental Advances 12, 100359.

Callahan, B.J., McMurdie, P.J., Rosen, M.J., Han, A.W., Johnson, A.J.A., Holmes, S.P., 2016. DADA2: High-resolution sample inference from Illumina amplicon data. Nature Methods 13, 581–583.

Camacho, C., Coulouris, G., Avagyan, V., Ma, N., Papadopoulos, J., Bealer, K., Madden, T.L., 2009. BLAST+: architecture and applications. BMC Bioinformatics 10, 1–9.

Cristina, A.M., Sara, F., Fausto, G., Vincenzo, P., Rocchina, S., Claudio, V., 2018. Degradation of post-consumer PLA: Hydrolysis of polymeric matrix and oligomers stabilization in aqueous phase. Journal of Polymers and the Environment 26, 4396–4404.

Danso, D., Schmeisser, C., Chow, J., Zimmermann, W., Wei, R., Leggewie, C., Li, X., Hazen, T., Streit, W.R., 2018. New insights into the function and global distribution of polyethylene terephthalate (PET)-degrading bacteria and enzymes in marine and terrestrial metagenomes. Applied and Environmental Microbiology 84, e02773–02717.

Das, S., Dash, H.R., 2014. Microbial bioremediation: A potential tool for restoration of contaminated areas, Microbial biodegradation and bioremediation. Elsevier, pp. 1–21.

de la Haba, R.R., Sánchez-Porro, C., Ventosa, A., 2011. Taxonomy, phylogeny, and biotechnological interest of the family Halomonadaceae, Halophiles and Hypersaline Environments: Current Research and Future Trends. Springer, pp. 27–64.

de Vogel, F.A., Schlundt, C., Stote, R.E., Ratto, J.A., Amaral-Zettler, L.A., 2021. Comparative genomics of marine bacteria from a historically defined plastic biodegradation consortium with the capacity to biodegrade polyhydroxyalkanoates. Microorganisms 9, 186.

Deepika, S., Jaya, M., 2015. Biodegradation of low density polyethylene by microorganisms from garbage soil. Journal of Experimental Biology and Agricultural Sciences 3, 1–5.

Delacuvellerie, A., Cyriaque, V., Gobert, S., Benali, S., Wattiez, R., 2019. The plastisphere in marine ecosystem hosts potential specific microbial degraders including *Alcanivorax borkumensis* as a key player for the low-density polyethylene degradation. Journal of Hazardous Materials 380, 120899.

Delaeter, C., Spilmont, N., Bouchet, V.M., Seuront, L., 2022. Plastic leachates: Bridging the gap between a conspicuous pollution and its pernicious effects on marine life. Science of the Total Environment 826, 154091.

Devi, R.S., Ramya, R., Kannan, K., Antony, A.R., Kannan, V.R., 2019. Investigation of biodegradation potentials of high density polyethylene degrading marine bacteria isolated from the coastal regions of Tamil Nadu, India. Marine Pollution Bulletin 138, 549–560.

Dintcheva, N.T., La Mantia, F., Stability, 2007. Durability of a starch-based biodegradable polymer. Polymer Degradation and Stability 92, 630–634.

Evtushenko, L.I., Takeuchi, M., 2006. The family microbacteriaceae, Springer, New York, NY.

Friedrich, J., Zalar, P., Mohorčič, M., Klun, U., Kržan, A., 2007. Ability of fungi to degrade synthetic polymer nylon-6. Chemosphere 67, 2089–2095.

Gambarini, V., Pantos, O., Kingsbury, J.M., Weaver, L., Handley, K.M., Lear, G., 2022. PlasticDB: a database of microorganisms and proteins linked to plastic biodegradation. Database, baac008.

Gardon, T., Huvet, A., Paul-Pont, I., Cassone, A.-L., Koua, M.S., Soyez, C., Jezequel, R., Receveur, J., Le Moullac, G., 2020. Toxic effects of leachates from plastic pearl-farming gear on embryo-larval development in the pearl oyster *Pinctada margaritifera*. Water Research 179, 115890.

Giacomucci, L., Raddadi, N., Soccio, M., Lotti, N., Fava, F., 2019. Polyvinyl chloride biodegradation by *Pseudomonas citronellolis* and *Bacillus flexus*. New Biotechnology 52, 35–41.

Govarthanan, M., Khalifa, A.Y., Kamala-Kannan, S., Srinivasan, P., Selvankumar, T., Selvam, K., Kim, W., 2020. Significance of allochthonous brackish water *Halomonas* sp. on biodegradation of low and high molecular weight polycyclic aromatic hydrocarbons. Chemosphere 243, 125389.

Gregory, M.R., 2009. Environmental implications of plastic debris in marine settings—entanglement, ingestion, smothering, hangers-on, hitch-hiking and alien invasions. Philosophical Transactions of the Royal Society B: Biological Sciences 364, 2013–2025.

Hahladakis, J.N., Velis, C.A., Weber, R., Iacovidou, E., Purnell, P., 2018. An overview of chemical additives present in plastics: Migration, release, fate and environmental impact during their use, disposal and recycling. Journal of Hazardous Materials 344, 179–199.

Hajighasemi, M., Tchigvintsev, A., Nocek, B., Flick, R., Popovic, A., Hai, T., Khusnutdinova, A.N., Brown, G., Xu, X., Cui, H., 2018. Screening and characterization of novel polyesterases from environmental metagenomes with high hydrolytic activity against synthetic polyesters. Environmental Science & Technology 52, 12388–12401.

Holzapfel, W.H., Wood, B.J., 2014. Lactic acid bacteria: biodiversity and taxonomy. John Wiley & Sons.

Jeszeová, L., Puškárová, A., Bučková, M., Kraková, L., Grivalský, T., Danko, M., Mosnáčková, K., Chmela, Š., Pangallo, D., 2018. Microbial communities responsible for the degradation of poly (lactic acid)/poly (3-hydroxybutyrate) blend mulches in soil burial respirometric tests. World Journal of Microbiology and Biotechnology Advances 34, 1–12.

Kalinová, J.P., Tříska, J., Vrchotová, N., Novák, J., 2016. Uptake of caprolactam and its influence on growth and oxygen production of *Desmodesmus quadricauda* algae. Environmental Pollution 213, 518–523.

Kasuya, K., Ohura, T., Masuda, K., Doi, Y., 1999. Substrate and binding specificities of bacterial polyhydroxybutyrate depolymerases. International Journal of Biological Macromolecules 24, 329–336.

Khandare, S.D., Chaudhary, D.R., Jha, B., 2021. Marine bacterial biodegradation of low-density polyethylene (LDPE) plastic. Biodegradation 32, 127–143.

Kim, H.-W., Jo, J.H., Kim, Y.-B., Le, T.-K., Cho, C.-W., Yun, C.-H., Chi, W.S., Yeom, S.-J., 2021. Biodegradation of polystyrene by bacteria from the soil in common environments. Journal of Hazardous Materials 416, 126239.

Klaeger, F., Tagg, A.S., Otto, S., Bienmüller, M., Sartorius, I., Labrenz, M., 2019. Residual monomer content affects the interpretation of plastic degradation. Scientific Reports 9, 2120.

Koutny, M., Amato, P., Muchova, M., Ruzicka, J., Delort, A.-M., 2009. Soil bacterial strains able to grow on the surface of oxidized polyethylene film containing prooxidant additives. International Biodeterioration and Biodegradation 63, 354–357.

Kumari, A., Chaudhary, D.R., Jha, B., 2019. Destabilization of polyethylene and polyvinylchloride structure by marine bacterial strain. Environmental Science Pollution Research 26, 1507–1516.

Lear, G., Maday, S.D., Gambarini, V., Northcott, G., Abbel, R., Kingsbury, J., Weaver, L., Wallbank, J., Pantos, O., 2022. Microbial abilities to degrade global environmental plastic polymer waste are overstated. Environmental Research Letters 17, 043002.

Martin, M., 2011. Cutadapt removes adapter sequences from high-throughput sequencing reads. EMBnet.journal 17, 10.

Masaki, K., Kamini, N.R., Ikeda, H., Iefuji, H., 2005. Cutinase-like enzyme from the yeast Cryptococcus sp. strain S-2 hydrolyzes polylactic acid and other biodegradable plastics. Applied and Environmental Microbiology 71, 7548–7550.

McLaren, M.R., 2020. Silva SSU taxonomic training data formatted for DADA2 (Silva version 138), Zenodo, Geneva, Switzerland, p. 10.5281/zenodo.3731174.

Molitor, R., Bollinger, A., Kubicki, S., Loeschcke, A., Jaeger, K.E., Thies, S., 2020. Agar plate-based screening methods for the identification of polyester hydrolysis by *Pseudomonas* species. Microbial Biotechnology 13, 274–284.

Montazer, Z., Habibi-Najafi, M.B., Mohebbi, M., Oromiehei, A., 2018. Microbial degradation of UV-pretreated low-density polyethylene films by novel polyethylene-degrading bacteria isolated from plastic-dump soil. Journal of Polymers and the Environment 26, 3613–3625.

Nadeem, H., Alia, K.B., Muneer, F., Rasul, I., Siddique, M.H., Azeem, F., Zubair, M., 2021. Isolation and identification of low-density polyethylene degrading novel bacterial strains. Archives of Microbiology 203, 5417–5423.

Nair, N.R., Sekhar, V.C., Nampoothiri, K.M., 2016. Augmentation of a microbial consortium for enhanced polylactide (PLA) degradation. Indian Journal of Microbiology 56, 59–63.

Navarro, D., Chaduli, D., Taussac, S., Lesage-Meessen, L., Grisel, S., Haon, M., Callac, P., Courtecuisse, R., Decock, C., Dupont, J., 2021. Large-scale phenotyping of 1,000 fungal strains for the degradation of non-natural, industrial compounds. Communications Biology 4, 871.

Nguyen, H.T.H., Qi, P., Rostagno, M., Feteha, A., Miller, S.A., 2018. The quest for high glass transition temperature bioplastics. Journal of Materials Chemistry A 6, 9298–9331.

Ohta, T., Tani, A., Kimbara, K., Kawai, F., 2005. A novel nicotinoprotein aldehyde dehydrogenase involved in polyethylene glycol degradation. Applied Microbiology and Biotechnology 68, 639–646.

Parales, R., Parales, J., Pelletier, D., Ditty, J., 2008. Diversity of microbial toluene degradation pathways. Advances in Applied Microbiology 64, 1–73.

Piubeli, F., Grossman, M., Fantinatti-Garboggini, F., Durrant, L., 2012. Identification and characterization of aromatic degrading *Halomonas* in hypersaline produced water and COD reduction by bioremediation by the indigenous microbial population using nutrient addition. Chemical Engineering Transactions 27, 385–390.

R Core Team, 2020. R: A language and environment for statistical computing. R Foundation for Statistical Computing, Vienna, Austria: R Foundation for Statistical Computing.

Raghul, S., Bhat, S., Chandrasekaran, M., Francis, V., Thachil, E., Technology, 2014. Biodegradation of polyvinyl alcohol-low linear density polyethylene-blended plastic film by consortium of marine benthic vibrios. International Journal of Environmental Sciences 11, 1827–1834.

Roman, J., McCarthy, J.J., 2010. The whale pump: marine mammals enhance primary productivity in a coastal basin. PloS One 5, e13255.

Romera-Castillo, C., Pinto, M., Langer, T.M., Álvarez-Salgado, X.A., Herndl, G.J., 2018. Dissolved organic carbon leaching from plastics stimulates microbial activity in the ocean. Nature Communications 9, 1430.

Sangale, M.K., Shahnawaz, M., Ade, A.B., 2019. Potential of fungi isolated from the dumping sites mangrove rhizosphere soil to degrade polythene. Scientific Reports 9, 5390.

Sarker, I., Moore, L.R., Paulsen, I.T., Tetu, S.G., 2020. Assessing the toxicity of leachates from weathered plastics on photosynthetic marine bacteria *Prochlorococcus*. Frontiers in Marine Science 7, 571929.

Sarkhel, R., Sengupta, S., Das, P., Bhowal, A., 2020. Comparative biodegradation study of polymer from plastic bottle waste using novel isolated bacteria and fungi from marine source. Journal of Polymer Research 27, 1–8.

Sekiguchi, T., Saika, A., Nomura, K., Watanabe, T., Watanabe, T., Fujimoto, Y., Enoki, M., Sato, T., Kato, C., Kanehiro, H., 2011. Biodegradation of aliphatic polyesters soaked in deep seawaters and isolation of poly (ɛ-caprolactone)-degrading bacteria. Polymer Degradation Stability 96, 1397–1403.

Shahnawaz, M., Sangale, M.K., Ade, A.B., 2016a. Bacteria-based polythene degradation products: GC-MS analysis and toxicity testing. Environmental Science and Pollution Research 23, 10733–10741.

Shahnawaz, M., Sangale, M.K., Ade, A.B., 2016b. Rhizosphere of *Avicennia marina* (Forsk.) Vierh. as a landmark for polythene degrading bacteria. Environmental Science and Pollution Research 23, 14621–14635.

Stringer, R., Johnston, P., 2001. Chlorine and the environment: an overview of the chlorine industry. Springer Science & Business Media.

Sudhakar, M., Doble, M., Murthy, P.S., Venkatesan, R., 2008. Marine microbe-mediated biodegradation of low-and high-density polyethylenes. International Biodeterioration and Biodegradation 61, 203–213.

Suzuki, M., Tachibana, Y., Kazahaya, J.-i., Takizawa, R., Muroi, F., Kasuya, K.-i., 2017. Difference in environmental degradability between poly (ethylene succinate) and poly (3-hydroxybutyrate). Journal of Polymer Research 24, 1–8.

Tachibana, K., Hashimoto, K., Yoshikawa, M., Okawa, H., 2010. Isolation and characterization of microorganisms degrading nylon 4 in the composted soil. Polymer Degradation and Stability 95, 912–917.

Takeyama, N., Huang, M., Sato, K., Galipon, J., Arakawa, K.J.M.R.A., 2020. Complete genome sequence of halomonas hydrothermalis strain slthf2, a halophilic bacterium isolated from a deep-sea hydrothermal-vent environment. 9, 10.1128/mra. 00294-00220.

Tetu, S.G., Sarker, I., Moore, L.R., 2020. How will marine plastic pollution affect bacterial primary producers? Communications Biology 3, 1–4.

Tetu, S.G., Sarker, I., Schrameyer, V., Pickford, R., Elbourne, L.D.H., Moore, L.R., Paulsen, I.T., 2019. Plastic leachates impair growth and oxygen production in *Prochlorococcus*, the ocean’s most abundant photosynthetic bacteria. Communications Biology 2, 184.

Vlaanderen, E.J., Ghaly, T.M., Moore, L.R., Focardi, A., Paulsen, I.T., Tetu, S.G., 2023. Plastic leachate exposure drives antibiotic resistance and virulence in marine bacterial communities. Environmental Pollution 327, 121558.

Wang, C., Chang, T., Yang, H., Cui, M.J.F.C., 2015. Antibacterial mechanism of lactic acid on physiological and morphological properties of *Salmonella enteritidis*, *Escherichia coli* and *Listeria monocytogenes*. Food Control 47, 231–236.

Xu, S., Yamaguchi, T., Osawa, S., Suye, S.-I., 2007. Biodegradation of poly (ε-caprolactone) film in the presence of Lysinibacillus sp. 70038 and characterization of the degraded film. Biocontrol Science 12, 119–122.

Yan, Z.-F., Wang, L., Xia, W., Liu, Z.-Z., Gu, L.-T., Wu, J., 2021. Synergistic biodegradation of poly (ethylene terephthalate) using *Microbacterium oleivorans* and *Thermobifida fusca* cutinase. Applied Microbiology and Biotechnology Advances 105, 4551–4560.

Zeng, X., Tang, W., Ye, G., Ouyang, T., Tian, L., Ni, Y., Li, P., 2010. Studies on disinfection mechanism of electrolyzed oxidizing water on *E. coli* and *Staphylococcus aureus*. Journal of Food Science 75, M253–M260.

Zhang, T., Cui, T., Cao, Y., Li, Y., Li, F., Zhu, D., Xing, J.J.A.v.L., 2022. Whole genome sequencing of the halophilic Halomonas qaidamensis XH36, a novel species strain with high ectoine production. 115, 545–559.

