## Supplementary materials for "An experimental approach to understanding plastic leachate impacts on marine microorganisms"

S1. Thermogravimetric analysis (TGA)

Both mechanically ground plastics and plastic tokens (harbouring marine plastic-associated microbial communities) were assessed using a Discovery TGA1-0329 Thermal Analysis instrument (TA Instruments, USA) to determine the inorganic matter concentrations for all polymer types. Samples (10 – 30 mg) were placed in 100 µl platinum high–temperature sample pans under the following programme: (i) heating ramp (10 °C/min) from ambient temperature to 800 °C under a nitrogen gas atmosphere (gas flow: 10 mL/min), and (ii) a constant temperature of 800 °C under an air atmosphere (gas flow: 10 mL/min) for 10 minutes. The inorganic content of plastics was then determined by the weight of residue remaining in each sample pan.

S2. Media

All glassware was first soaked in a 10% Trigene solution and rinsed with distilled water before autoclaving at 121°C for 20 mins. Glassware was then rinsed in acetone and sterile deionised water three times to remove any residues that might influence plastic weathering or leachate composition. Plastic powders were suspended in artificial seawater salts (ASWS) broth at a concentration of 100 g $L^{-1}$ (pH 7.5 ± 0.5 units). ASWS media comprised 24.6 g $L^{-1}$ NaCl, 0.67 g $L^{-1}$ KCl, 1.36 g $L^{-1}$ CaCl $2H_2O$, 6.29 g $L^{-1}$ $MgSO_4 \cdot 7H_2O$, 4.66 g $L^{-1}$ $MgCl \cdot 6H_2O$ and 0.18 g $L^{-1}$ $NaHCO_3$, to mimic the basal chemical constituents within the marine environment. ASWS was dissolved in Type I grade (Ultrapure) water and autoclaved at 120°C before plastic powder addition. Four 250 ml replicates of each plastic were thermally weathered for three months in the dark at 80°C, along with a set of control samples (ASWS lacking ground plastics). Samples were shaken periodically, once a week, to separate any plastic clumps. After weathering, leachates were filtered using a 40 µm steel filter that had been rinsed in acetone and sterile deionised water three times to remove larger plastic particles (Rummel et al. 2019) and then filtered using a 0.2 µm Nalgene sterile filter and a vacuum pump (Capolupo et al. 2020) to remove smaller particles, including microbial cells. The filtered leachates were adjusted to ~pH 7 using 5 M NaOH and stored in sterile glassware, wrapped in foil at 4°C until required.

S3. Biological contaminant testing

In triplicate, 100 µl of each filtered plastic leachate was spread onto nutrient agar and incubated at 25°C for seven days; no colonies formed on agar plates. Additionally, 250 µl of each leachate type was subjected to DNA extraction and PCR (using primers for 16S and ITS rRNA gene regions, described in the methods section on DNA extraction, metabarcoding and amplicon sequencing); no samples yielded any PCR product, as evidenced by gel electrophoresis. Therefore, there was no evidence of viable bacteria or bacterial DNA in the leachates.

S4. Marine biofilm community sampling

To isolate marine plastic-associated microbial communities, we installed LLDPE, PA6, PET and PLA plastic tokens (Figure S2) in the Lighter Basin Marina of Tāmaki Makaurau-Auckland's Viaduct Basin, Aotearoa-New Zealand at latitude 36°50'42.4''S and longitude 174°45'29.6''E (Figure S2C). Paddles from each of the four plastic types were installed at depths between 20 to 60 cm from the ocean surface on stainless-steel (316) poles using a stainless-steel (316) frame attached to a pontoon (Figure S2A) on 17$^{th}$ March 2021 during Austral summer (Laroche et al. 2023). Two paddles (75 mm x 50 mm x 3 mm) of each plastic type were excised from the structure after two weeks of incubation on 31$^{st}$ March 2021. During excision, sterile Fisherbrand sample bags (CAT No. 14955189; Thermo Fisher Scientific, Waltham, MA, USA) were placed over each paddle before they were cut from the structure using pipe cutters to remove the integrated arm (Figure S2B). Paddles were transported on ice to the lab, where biofilms were scraped off each paddle using a flat-edged razor blade, and each sample was stored separately in 15% glycerol stock solutions at -80°C, i.e. two replicates of each plastic type for a total of eight samples, stored until required.

S5. Quantifying cell numbers

During the preconditioning period, cell counts were estimated via optical density measures ($OD_{600nm}$) using a SpectraMax ID3 plate reader (Molecular Devices, LLC, San Jose, California, U.S.A), and use of a LIVE/DEAD *Bac*Light Bacterial Viability Kit (ThermoFisher Scientific, Waltham, MA, U.S.A). Cells were viewed using a Leica DMR microscope at 63x magnification using an oil immersion lens under green fluorescence protein (GFP), and tetramethylrhodamine (TRITC) filters for SYTO 9 and propidium iodine imaging, respectively. Images were captured using a Jenoptik Gryphax Kapella camera (Jenoptik, Jena, Germany). Estimates of percentage cell viability were calculated by counting the number of green fluorescent cells divided by the total number of cells. Cell counts were automated using a 'Biofilm Viability Checker' macro in ImageJ (Mountcastle et al. 2021) with default parameters, except for adjusted image gamma values.

S6. Analysis of GC-MS data

The organic chemical composition of each plastic leachate type was assessed using a non-targeted approach by solvent extraction gas chromatography-mass spectrometry (GC-MS). Supernatants were concentrated using a SpeedVac High-Capacity Vacuum Concentrator (Thermo Fisher Scientific, Waltham, MA, USA) to roughly 5 ml over 24 hours. In addition to these samples, both MilliQ Water and Ethyl Acetate were introduced in triplicate as GC-MS controls. Samples were processed by adding a 4 ml volume of ethyl acetate, laced with surrogate internal standards (i.e., 25.08 ng naphthalene-*d*8, 4.86 ng chrysene-*d*12) to 4 ml of each of the concentrated samples, and then sonicated at 40 KHz for 30 min at 65°C. The entire solvent layer was transferred to a clean test tube using a glass Pasteur pipette. Solvent layers were concentrated by adding (~50 mg) anhydrous $Na_2SO_4$ to each sample tube and evaporating solvents under a gentle stream of $N_2$ to approximately 500 µl before being transferred into a GC vial and analysed via GC-MS (Agilent 7890A GC equipped with an Agilent 5975C Mass Selective Detector).

Using Agilent Chemstation software (Agilent Technologies, Santa Clara, CA 95051, USA), individual sample chromatogram peaks were integrated using a non-redundant National Institute of Standards and Technology (NIST) 2017 library database (NIST 2017), taking the top 10 hits with a minimum purity of 50 to build a subset library database. Chromatograms were then batch peak deconvoluted and identified using the custom NIST17 database and an R package for untargeted metabolomic batch processing (Guo et al. 2020), with match factors of 45%. Batch integration and library contamination analyses were processed following the authors' instruction (Guo et al. 2020) before cumulative sum scaling (css) correction and normalisation. Outputs were processed in R studio (R Core Team, 2020) using the 'vegan' R package (version 4.6-4; Oksanen, 2022) to produce NMDS plots. Data were statistically tested using permutational multivariate analysis of variance (PERMANOVA; Anderson, 2008) with 999 permutations of the data, carried out on Bray-Curtis dissimilarity matrices using the vegan 'adonis' function. Differential abundance analysis assessed the data distribution using the normality Shapiro and Kolmogrov-Smirov tests, followed by the Kruskal Wallis test for differential abundance, available in base R packages (R Core Team, 2020). Within base R, Dunn's test for differential abundance compared samples representing the same treatment condition to identify organic components specific to carbon media conditions.

S7. DNA extraction, metabarcoding and amplicon sequencing

Microbial DNA was extracted from the centrifuged pellets of each sample using a DNeasy PowerSoil Pro kit (Qiagen, Hilden, Germany) following the manufacturer's instructions, except for the mechanical lysis step, which was performed using a TissueLyser II (Cat No. 85300; Qiagen, Hilden, Germany) for 2 mins at 30 Hz. For samples with no visible pellet, ~200 µl of the supernatant was used; this was required for several LLDPE and PET 0% carbon samples. DNA was eluted in 100 µl 10 nM Tris Buffer. DNA concentrations were assessed using a Nanodrop photometer (Implen Nanophotometer, Munich, Germany) and stored at -20°C until required. Following PCR, agarose gel electrophoresis was performed to verify the amplification of microbial DNA. A ZymoBIOMICS®

Microbial Community Standard was included in PCR amplifications of 16S rRNA gene regions and subsequently sequenced to check for biased amplification. A Qubit double-stranded DNA (dsDNA) High Sensitivity Assay Kit (Invitrogen, Thermo Fisher Scientific, Waltham, MA, USA) was used to quantify the DNA concentration before and after DNA purification. DNA was purified using a DNA clean and concentrator-5 kit (Zymo Research, Irvine, CA, USA) and eluted in 12 µl DNA elution buffer, according to manufacturer instructions. The DNA concentration in each sample ranged from 110 µg to 250 µg (per the manufacturer's guidelines). The Auckland Genomics facility (The University of Auckland, A-NZ) attached Nextera XT dual indices to amplicons (Illumina Inc, San Diego, CA, USA) and conducted DNA sequencing on an Illumina MiSeq instrument using 2-by-300-bp V3 chemistry. The mock community standard contained 23,542 filtered sequences, above the average of 19,655 for all other samples. All taxa were identified to at least the genus level per the ZymoBIOMICS® Microbial Community Standard taxa list for 16S rRNA gene sequencing, with no evidence of contamination. There was, however, evidence of biases during either PCR and/or sequencing, with *Limosilactobacillus* sp. dominating the community (Figure S5). Due to the limited utility of the mock community sample for the analysis of fungal data, containing only two species, it was not sequenced.

S8. RNA extraction and sequencing

Microbial RNA was extracted from 20 ml pelleted samples using an RNeasy PowerBiofilm kit (Qiagen, Hilden, Germany) following the manufacturer's instructions, except for the mechanical lysis step, which was performed using a TissueLyser II (Cat No. 85300; Qiagen, Hilden, Germany) for 2 mins at 30 Hz. As per DNA extractions, for samples that fell below pellet weight guidelines (110 - 200 µg), up to 200 µl of supernatant was used. RNA was eluted in 100 µl RNase-free water (Qiagen, Hilden, Germany). Per the manufacturer's instructions, DNA was removed using a TURBO DNA-free™ kit (Life Technologies Cat No. AM1907, Thermo Fisher Scientific, USA). The absence of DNA was verified via long cycle (35 cycles) 16S rRNA gene PCR amplification with the universal amplicon primer pair 341F and 785R and visualisation via gel electrophoresis. DNA-free RNA was cleaned and concentrated with an RNA Clean and Concentrator-5 (Cat. No. R1013, Zymo Research, Irvine, CA, USA). A260/230 sample purity was assessed using a nanophotometer (Implen, Munich, Germany). RNA concentration was evaluated using a Qubit™ RNA high-sensitivity Assay kit (Cat. No. Q32855, Life Technologies, Thermo Fisher Scientific, OR, USA) and Qubit™ 3 fluorometer, following the manufacturer's instructions. Finally, RNA quality was assessed with an Agilent RNA 6000 Nano Kit and Bioanalyzer 2100 following the manufacturer's instructions.

S9. DNA amplicon bioinformatics and quantitative analysis

All quantitative and statistical analyses and data visualisation were performed using R (version 4.2.2) (R Core Team, 2020). Rarefaction curves of ASV data were plotted (Figure S6) to evaluate if sufficient DNA sequencing depth had been achieved. To achieve standard sequencing depth and comparability of diversity, samples were rarefied using the ‘vegan’ R package (version 4.6-4; Oksanen, 2022). Neither bacterial nor fungal ASV tables contained ASVs not assigned to the kingdom ‘Bacteria’ or ‘Fungi’, respectively. Samples containing less than 1,000 ASVs were removed. ASV tables were then randomly subsampled to the minimum non-chimeric library size (rarefying; set.seed(123), bacteria n = 1151, fungi n = 1469). ASV rarefaction curves indicate sufficient sequencing depth to represent community characteristics for ITS-sequenced samples, whereas further sequencing could be desirable for 16S rRNA samples.

Permutational multivariate analysis of variance (PERMANOVA; Anderson, 2008) with 999 permutations was carried out on Bray-Curtis dissimilarity matrices using the vegan ‘adonis’ function to evaluate whether plastic leachate exposure had a significant effect ($P < 0.05$) on the composition of bacterial communities. PERMANOVAs were not carried out on pooled fungal community data due to insufficient samples. Pairwise PERMANOVA with the ‘pairwise.adonis’ function was conducted to compare the composition of bacteria in different plastic leachate-exposed communities. Indicator value (IndVal) analysis (with 999 permutations and $P < 0.05$) was performed to evaluate which bacterial ASVs contributed most to the observed differences in community composition (Dufrêne and Legendre 1997). Previously reported fungal degraders were too low in abundance to be considered indicator species. The relative abundances of bacterial indicator genera were plotted in relation to plastic leachate media type, also plotting their respective sensitivity and specificity values from IndVal analyses.

Two-dimensional non-metric multidimensional scaling (nMDS) ordinations were plotted using ‘ggplot2’ based on microbial community Bray-Curtis dissimilarities (using rarefied ASVs). Ellipses were drawn around data points using the ‘ggforce::geom_mark_ellipse’ inside the ‘ggforce’ package for ggplot2, assuming a multivariate t-distribution. nMDS plots were plotted based on the plastic leachate media type and percentage of media carbon, with phyla vectors added as a separate plot using the ‘envfit’ package.

### S10. Meta-transcriptome assembly and analysis

All PLA and control (ASWS and precondition) transcriptomic sequences had Nextera adapters and low-quality reads (phred score $<30$) trimmed using Trimmomatic (version 0.39) (Bolger et al. 2014). Sequences were removed if they had a $>2$ bp mismatch, paired read score of 30, or single read score of 10, and leading and tailing ends had a quality N score $<3$. Trimmed read quality was assessed using

FastQC v0.11.7 (Andrews, 2010). Ribosomal RNA was removed from the dataset using sortmeRNA (version 4.3.6) (Kopylova et al. 2012), using a non-redundant silva database (Pruesse et al. 2007), and Rfam database (Kalvari et al. 2021) (accessed 1st April, 2023). The remaining reads were *de novo* co-assembled using rnaSPAdes with the default kmer size range (Bushmanova et al. 2019). Assembly quality was assessed using the pseudoalignment tool Kallisto (Bray et al. 2016) and by the presence of universal coding regions using BUSCO (Simão et al. 2015). Transcripts were then clustered based on 95% confidence matching isoforms to reduce redundancy in sequence analysis. This was achieved using CD-HIT (Fu et al. 2012). Following clustering, the metatranscriptome was annotated by searching using DIAMOND blastp (Buchfink et al. 2021) against a curated SWISS-PROT/UniProt database (Boeckmann et al. 2003), accessed in May 2023, for homology assignments. Finally, transcripts were searched using DIAMOND blastp against a plastic-degrading protein database, PlasticDB (Gambarini et al., 2022).

Of 124 bacterial BUSCO sequences, the transcriptome assembly contained 72.6% of these as complete sequences (13.7% single, 58.9% duplicated) and a total of 714,004 coding sequence regions. A total of 1990 of these putative coding region transcripts were aligned with 161 unique previously reported putative plastic degradation encoding genes identified by the PlasticDB (Gambarini et al. 2022). For differential expression analysis, trimmed and filtered transcripts were aligned to the assembled metatranscriptome to estimate transcript abundance using HISAT2 (Kim et al. 2019). Sequence Alignment/Map (SAM) files were compressed to binary format (BAM) using SAMtools (version 1.15.1 (Li et al. 2009)) before transcriptomic features were counted using the featureCount function of Subread (version 2.0.3, (Liao et al. 2014)). The average read mapping success rate was 94.73%. Transcript count data were then processed in R (R Core Team 2020) using three packages for differential expression analysis: DESeq2 (Love et al. 2014), EdgeR (Robinson et al. 2010) and Limma + voom (Law et al. 2014), with results compared downstream to capture the breadth of differentially abundant transcripts. Differentially expressed transcripts with an FDR $P$ value > 0.05 were removed from subsequent analyses. Analyses were visualised in R with ggplot2 (Wickham and Wickham 2016).

S11. Taxonomic variation in fungal community composition across leachate type and carbon concentration

Fewer samples contained sufficient fungal ASVs for analysis compared to bacterial samples post-filtering (samples removed included PLA-50 and PLA-75, which had 342 and 39 sequences, respectively). ASVs were identified as belonging to the phylum *Basidiomycota* or *Ascomycota*. These ASVs were all characterised to at least the family level, but only some to the genus level.

*Trichosporonaceae* dominated all pooled samples, including the initial preconditioned community; LLDPE-0 and PA-0 had no other ASVs identified within the community (Figure S11). The precondition community had the greatest number of unique ASVs, followed by PLA-0. These two communities only shared two ASVs, identified as *Trichosporonaceae* and *Metschnikowiaceae*. PLA-0 was the only community to contain *Psathyrellaceae*, making up 11.6% of the community, whereas PET-75 was the only plastic leachate type to have *Coniochaeraceae,* making up 22.3% of the overall community composition (Figure S11). All other ASVs were <5% of their respective community, except for ASV 22, identified as *Xylariales Incertae sedis*, making up 11.3% of the preconditioned community (Figure S11).

### Supplementary Figures

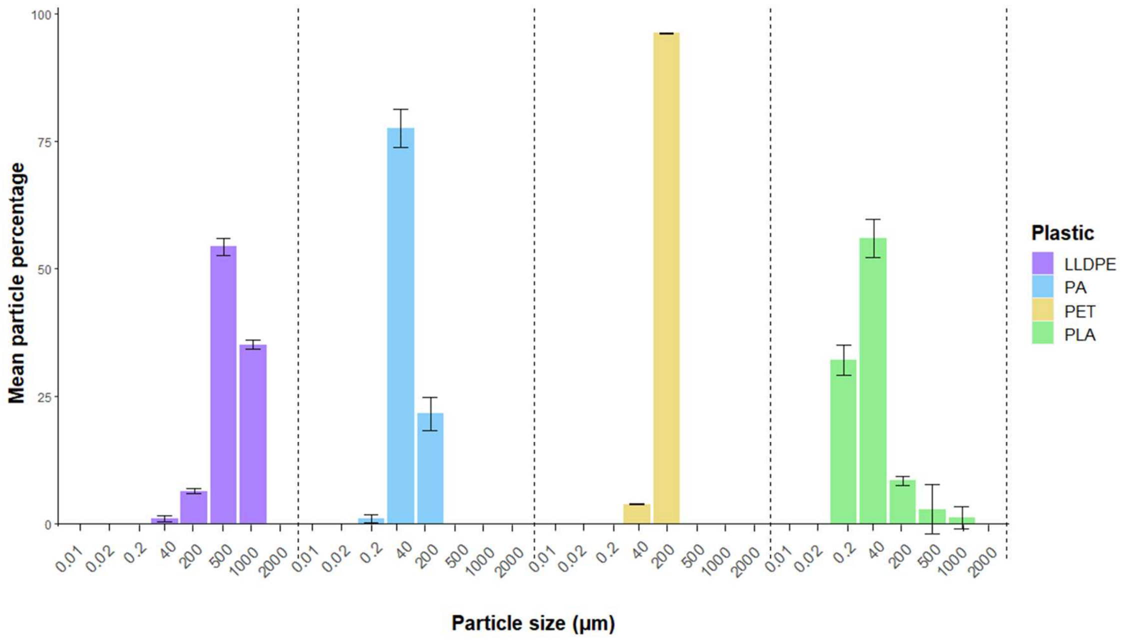


**Figure S1**. Mean percentage of mechanically ground microplastic fragments in different size groups used for artificial weathering and leachate generation at eight size ranges (µm). Standard deviations from the mean are represented by error bars.

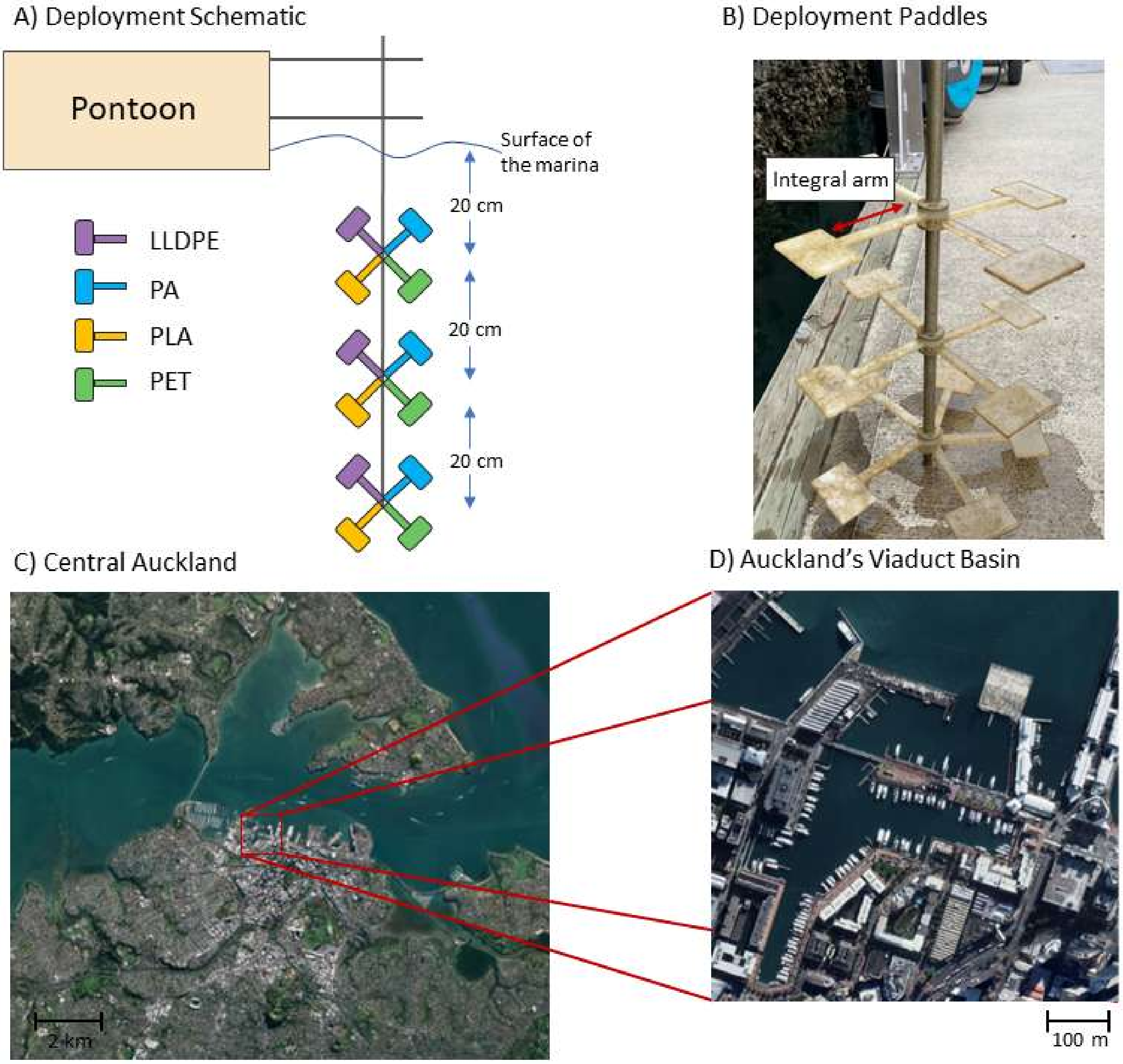


**Figure S2.** Stainless steel deployment structure affixed to a pontoon in the Central Auckland Region, Aotearoa-New Zealand. (A) Deployment schematic; (B) Plastic paddles attached to the deployment pole (out of water), indicating the integral arm; (C) Study location in the Central Auckland Region of Auckland, New Zealand, with inset (D) providing a closer view of Auckland's Viaduct Marina.

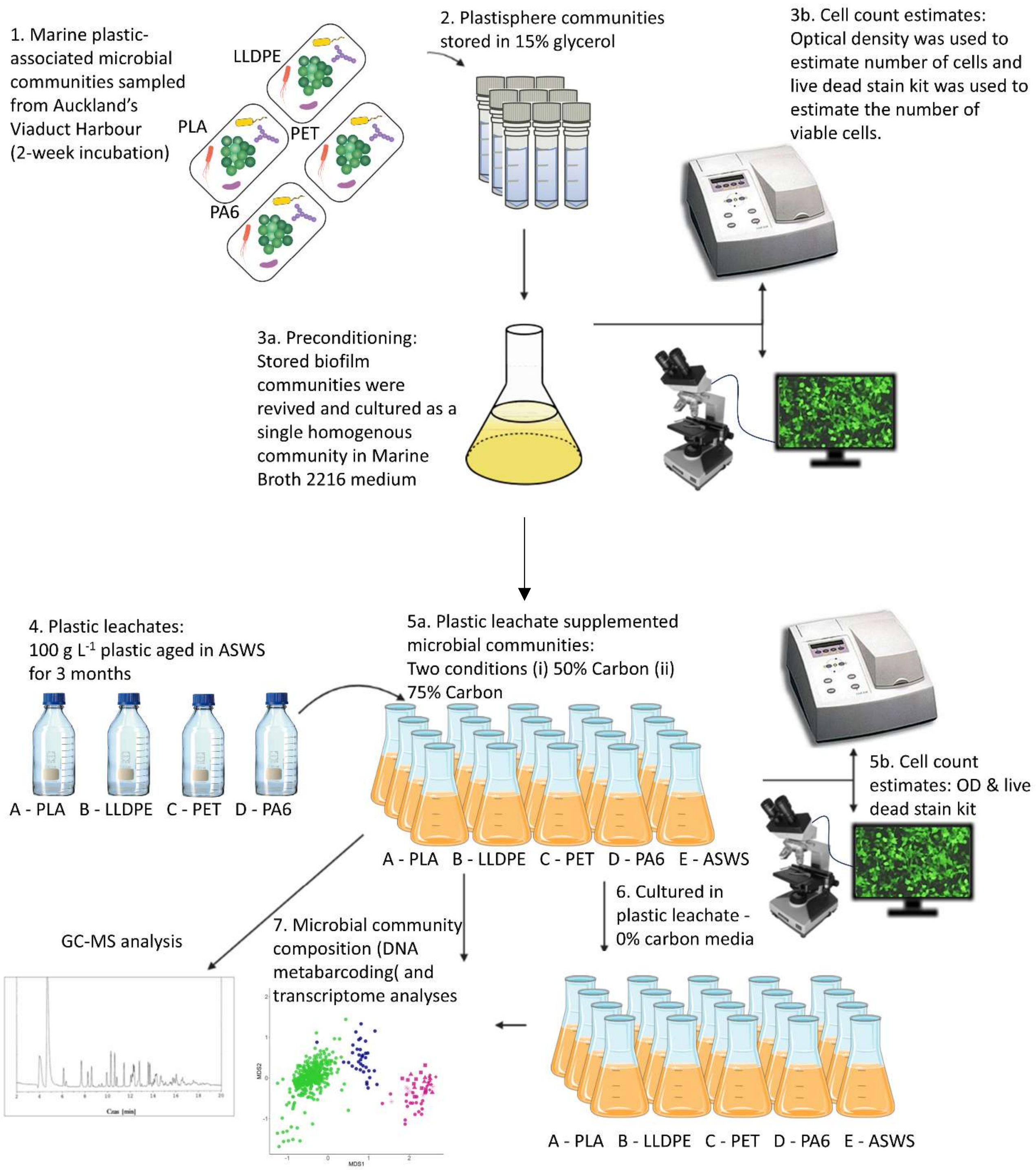


**Figure S3**. Schematic of the experimental design to investigate the impact of plastic leachates (from linear low-density polyethylene [LLDPE], polyamide-6 [PA6], polyethylene terephthalate [PET], polylactic acid [PLA]) and a control sample (artificial seawater salt broth [ASWS]) on plastic-derived marine microorganisms.

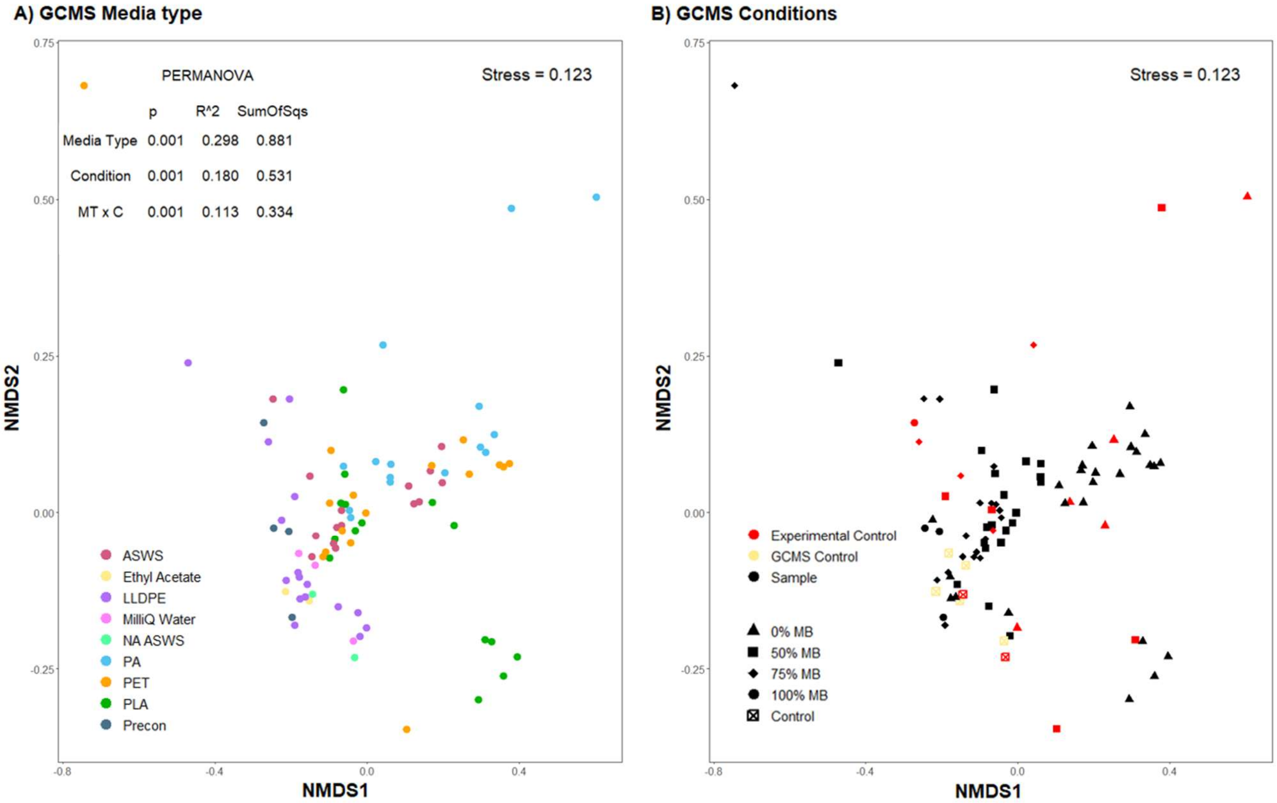


**Figure S4.** Plastic leachate composition two weeks post incubation as determined by GCMS. Non-metric multidimensional scaling (NMDS) ordination of Bray-Curtis similarities for (A) plastic leachate data grouped by leachate type ((ASWS) artificial seawater salt, (LLPDE) linear low-density polyethylene, (PA) polyamide-6, (PET) polyethylene terephthalate, (PLA) polylactic acid, (PreCon) the preconditioned microbial community control group, (ethyl acetate) a group of GCMS controls including; ethyl acetate + internal standard (naphthalene – d8), MilliQ Water, and (NA ASWS) a non-aged ASWS control). (B) Microbial community supernatant data stratified by growth media condition (0%, 50% or 75% carbon), with both incubated experimental controls and non-incubated (GCMS controls) gas chromatography–mass spectrophotometry controls. GC-MS controls were excluded from PERMANOVA analyses.

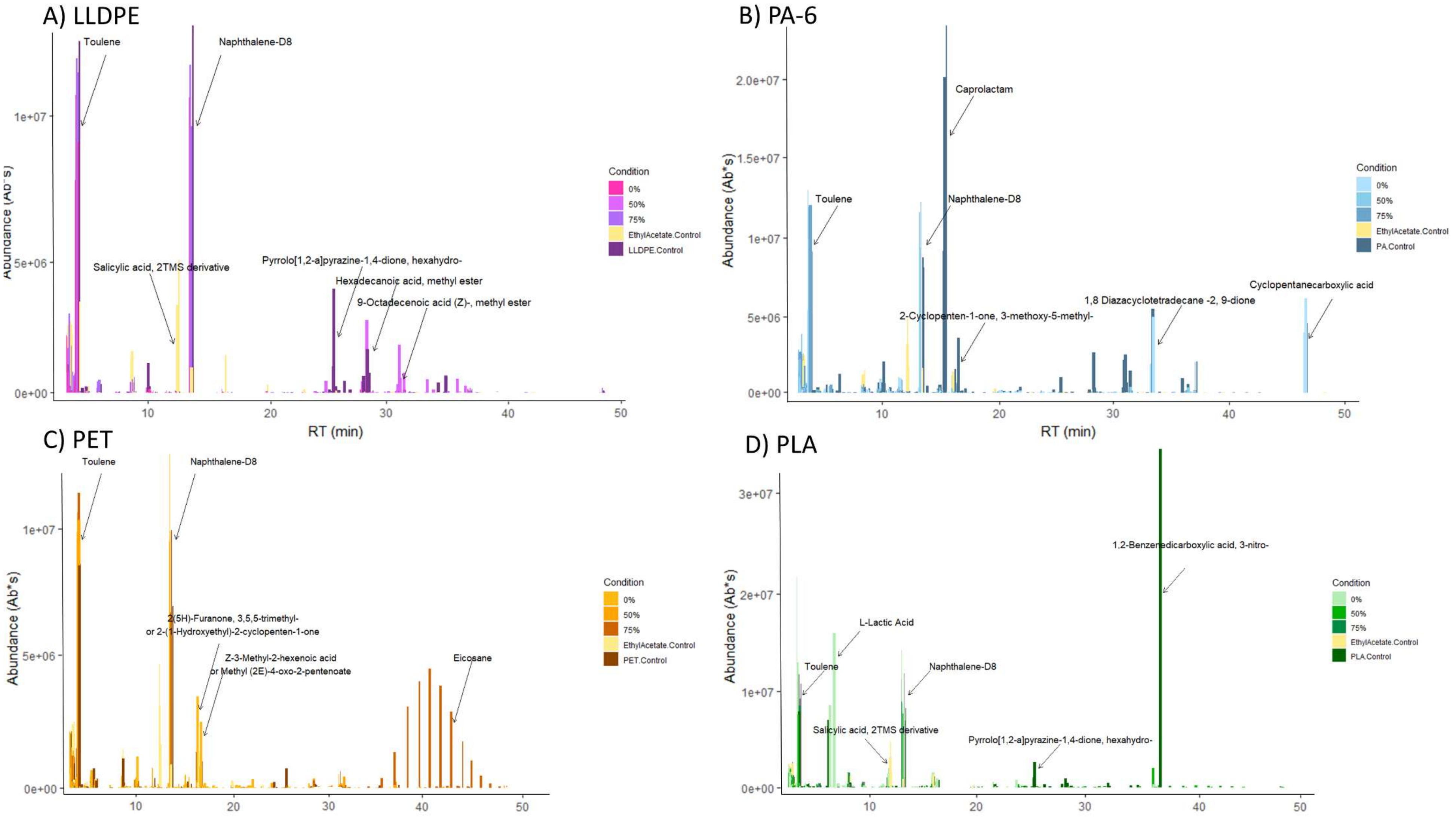


**Figure S5.** Organic components abundance based on the absorption peak surface area of individual sample chromatograms identified by a non-redundant NIST17 (National Institute of Standards and Technology) database. (a) Linear Low-density polyethylene (LLDPE), (b) Polyamide-6 (PA6), (c) Polyethylene terephthalate (PET), and (d) Polylactic acid (PLA) Samples are separated by percentage media carbon; 0%, 50%, 75% and both experimental and Gas Chromatography-Mass Spectrophotometry (GCMS) controls.

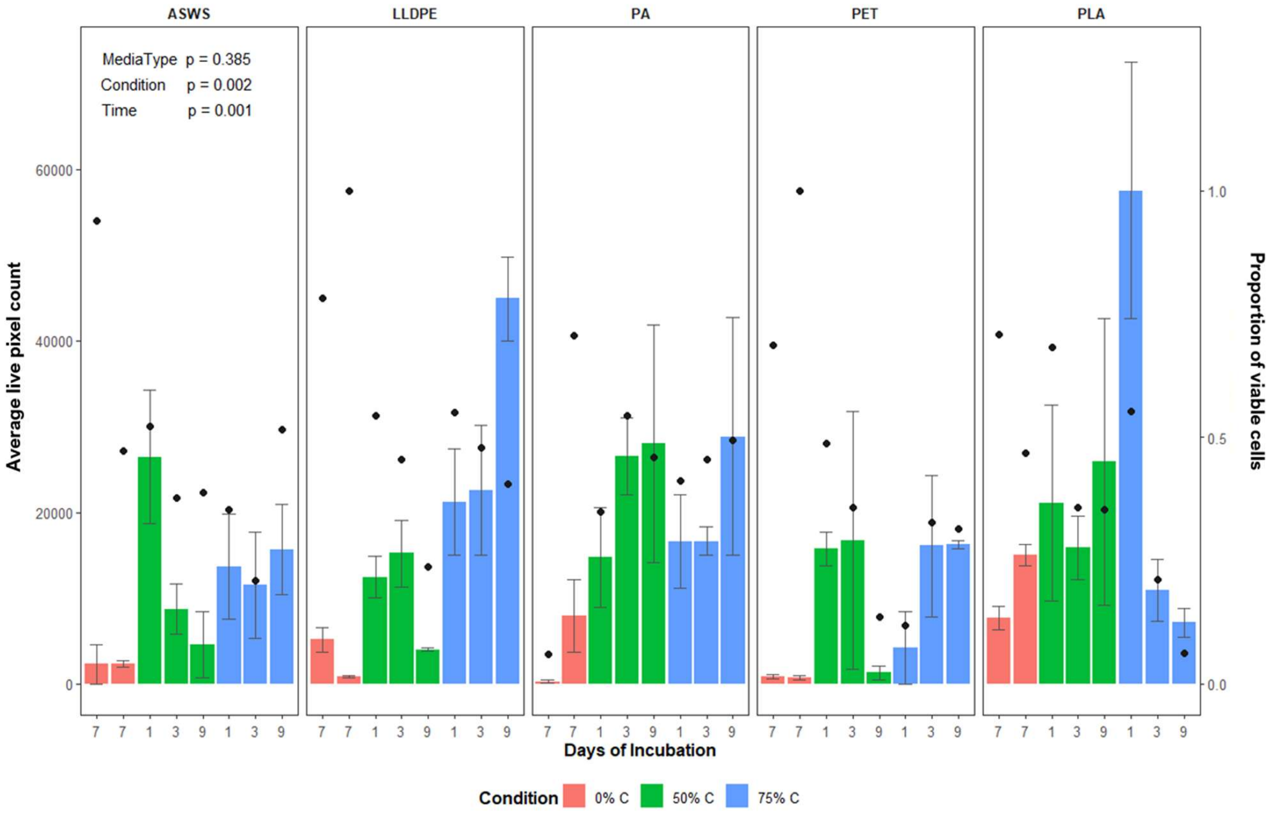


**Figure S6.** Average cell viability estimates based on SYTO9 and propidium iodide fluorescence microscopy pixel counts. Average pixel counts are shown as bars with standard deviation denoted as error bars. The proportion of total viable cells is shown as dots. Total pixels were counted using a modified ImageJ macro (Schneider et al. 2012), and the percentage of viable cells was estimated using a biofilm viability checker (Mountcastle et al. 2021).

| | Day 1 | Day 3 | Day 9 | 0% Carbon |
|---|---|---|---|---|
| ASWS | 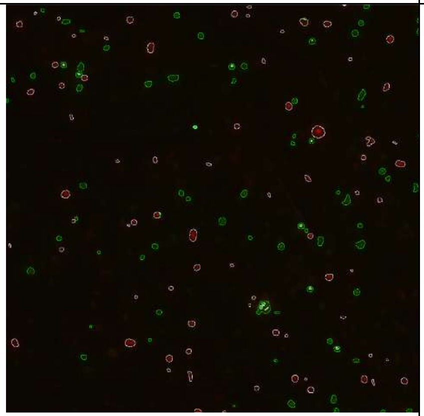 | 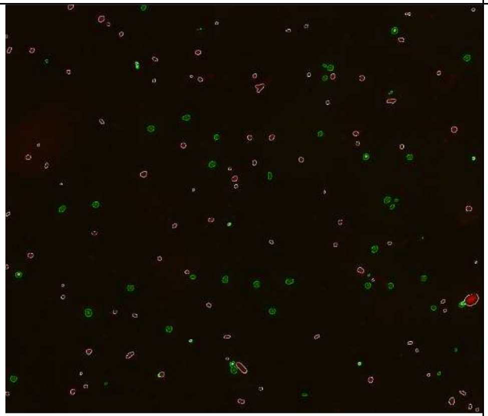 | 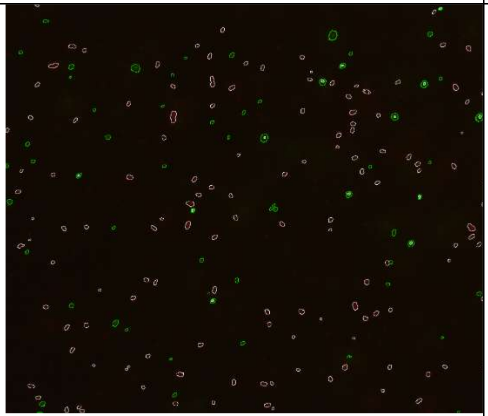 | 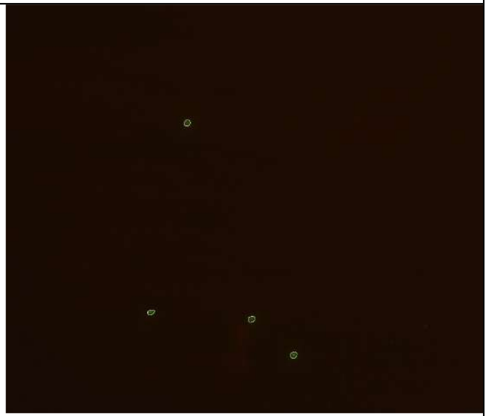 |
| LLDPE | 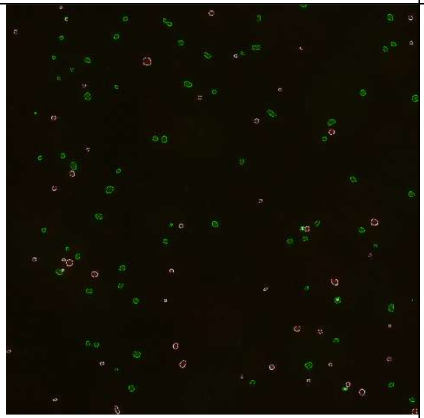 | 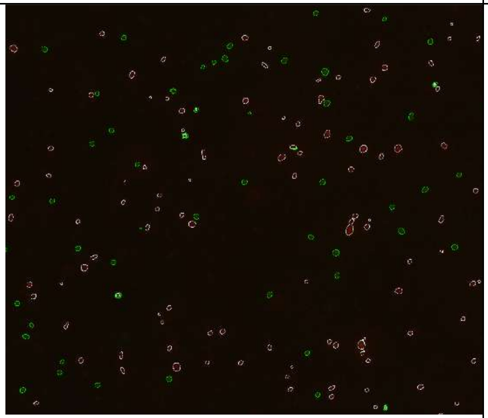 | 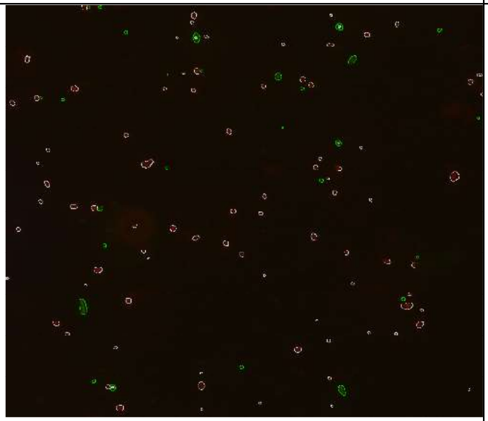 | 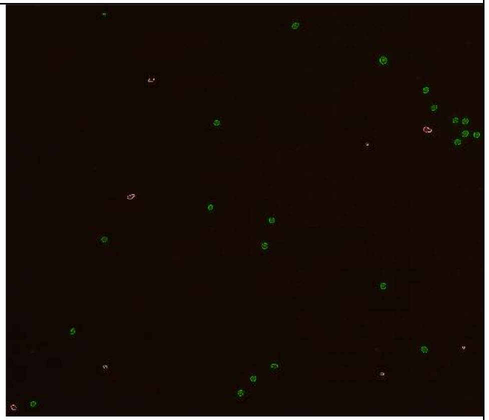 |

| | | | | |
|---|---|---|---|---|
| PA | 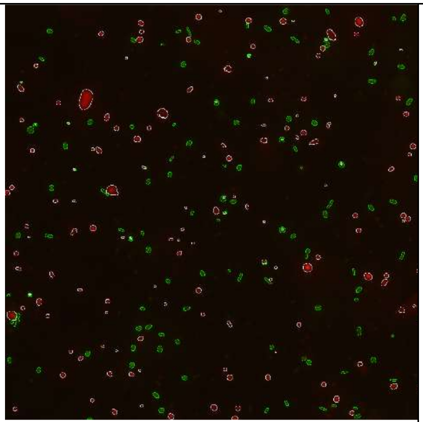 | 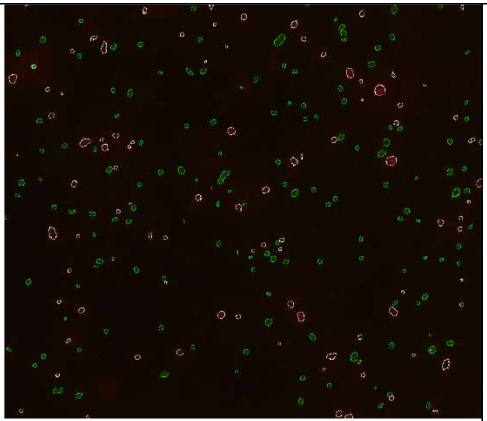 | 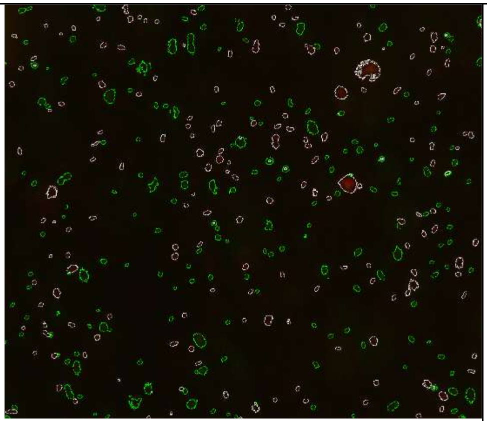 | 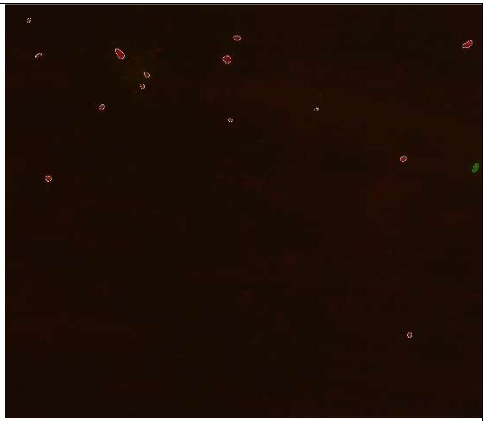 |
| PET | 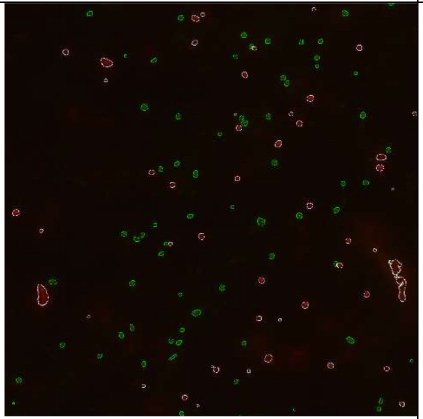 | 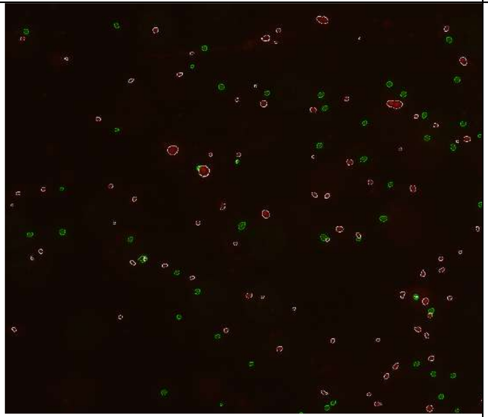 | 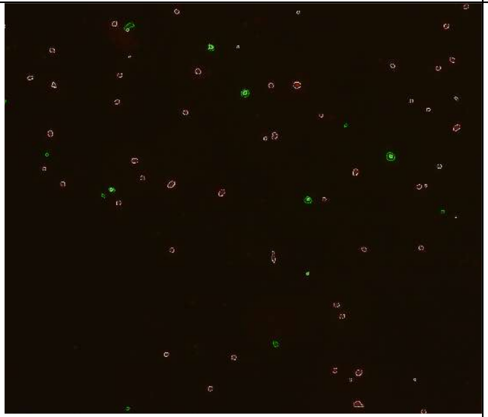 | 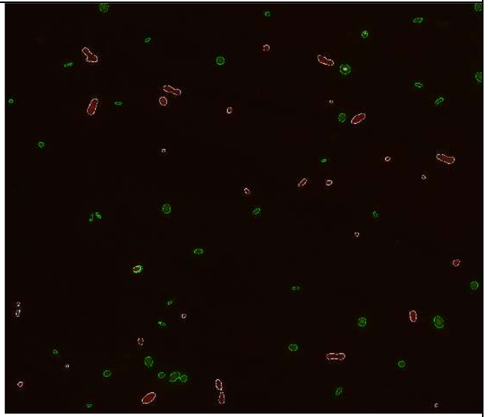 |

| | | | | |
|---|---|---|---|---|
| PLA | 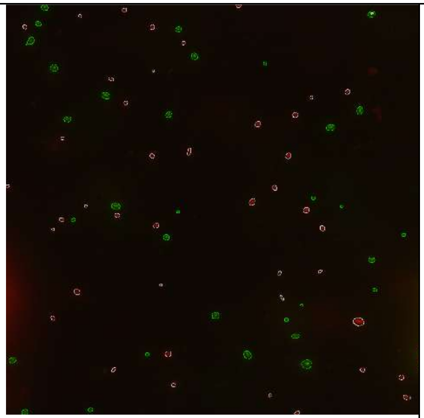 | 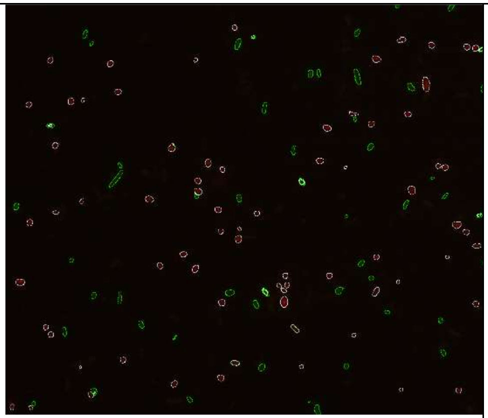 | 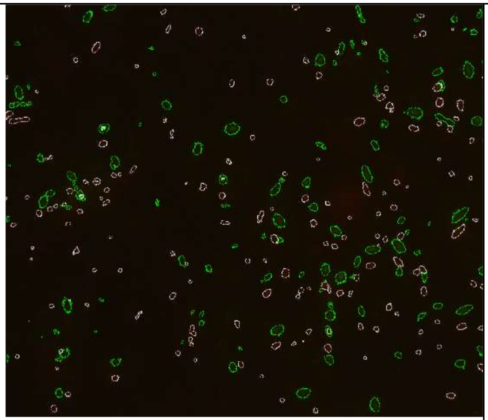 | 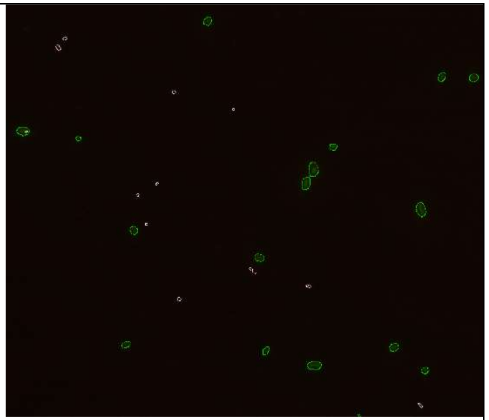 |
| Precon ditione d | 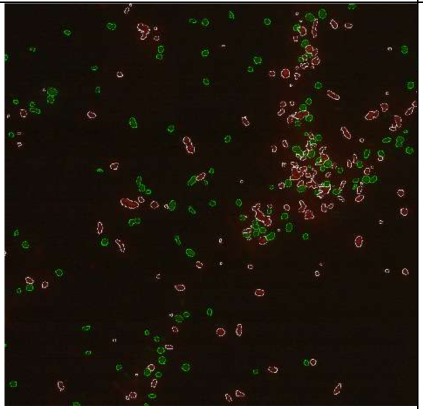 | 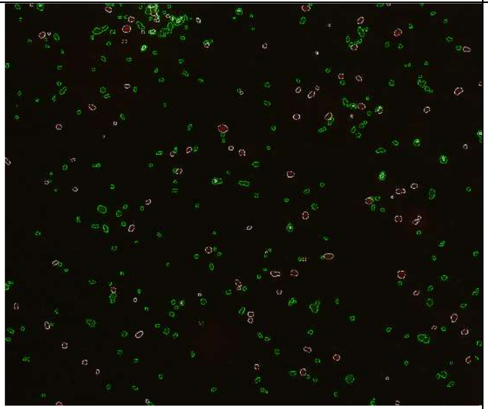 | 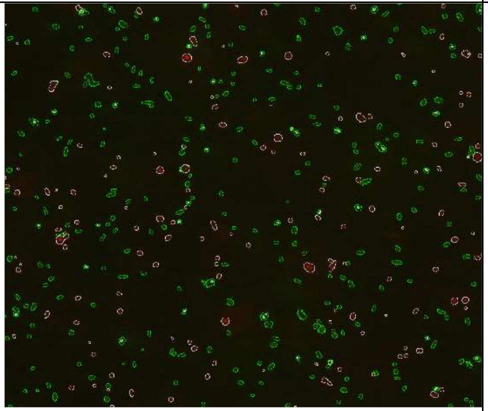 | |

**Figure S7.** Example images of cells stained with either SYTO9 or propidium iodine after ImageJ macro cell count (National Institutes of Health, Bethesda, Maryland, U.S.A) (Mountcastle et al. 2021). Cells circled in white indicate dead/dying cells; cells circled in green indicate live cells.

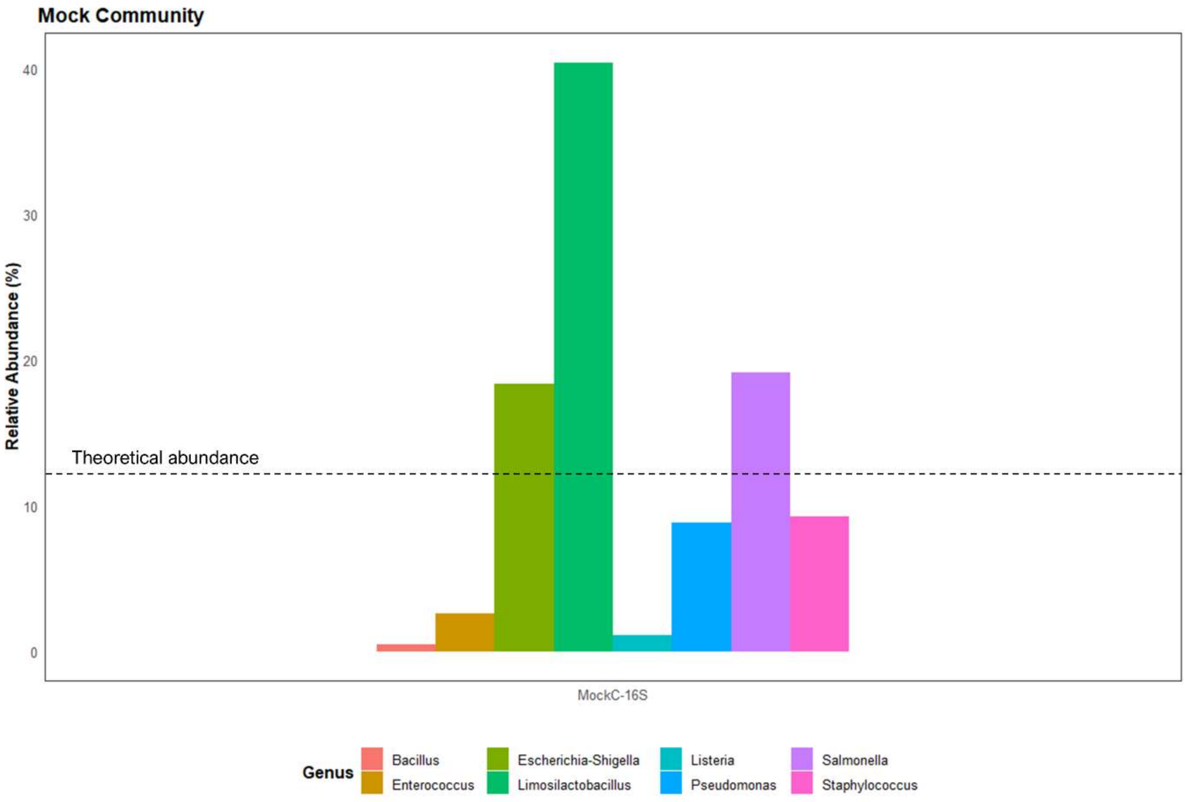


**Figure S8.** Relative abundance following sequencing and read processing of mock community standard (ZymoBIOMICS® Microbial Community Standard) for 16S rRNA reads. Theoretical composition based on genomic bacterial DNA: *Listeria monocytogenes* - 12%, *Pseudomonas aeruginosa* - 12%, *Bacillus subtilis* - 12%, *Escherichia coli* - 12%, *Salmonella enterica* - 12%, *Lactobacillus fermentum* - 12%, *Enterococcus faecalis* - 12%, and *Staphylococcus aureus* - 12%.

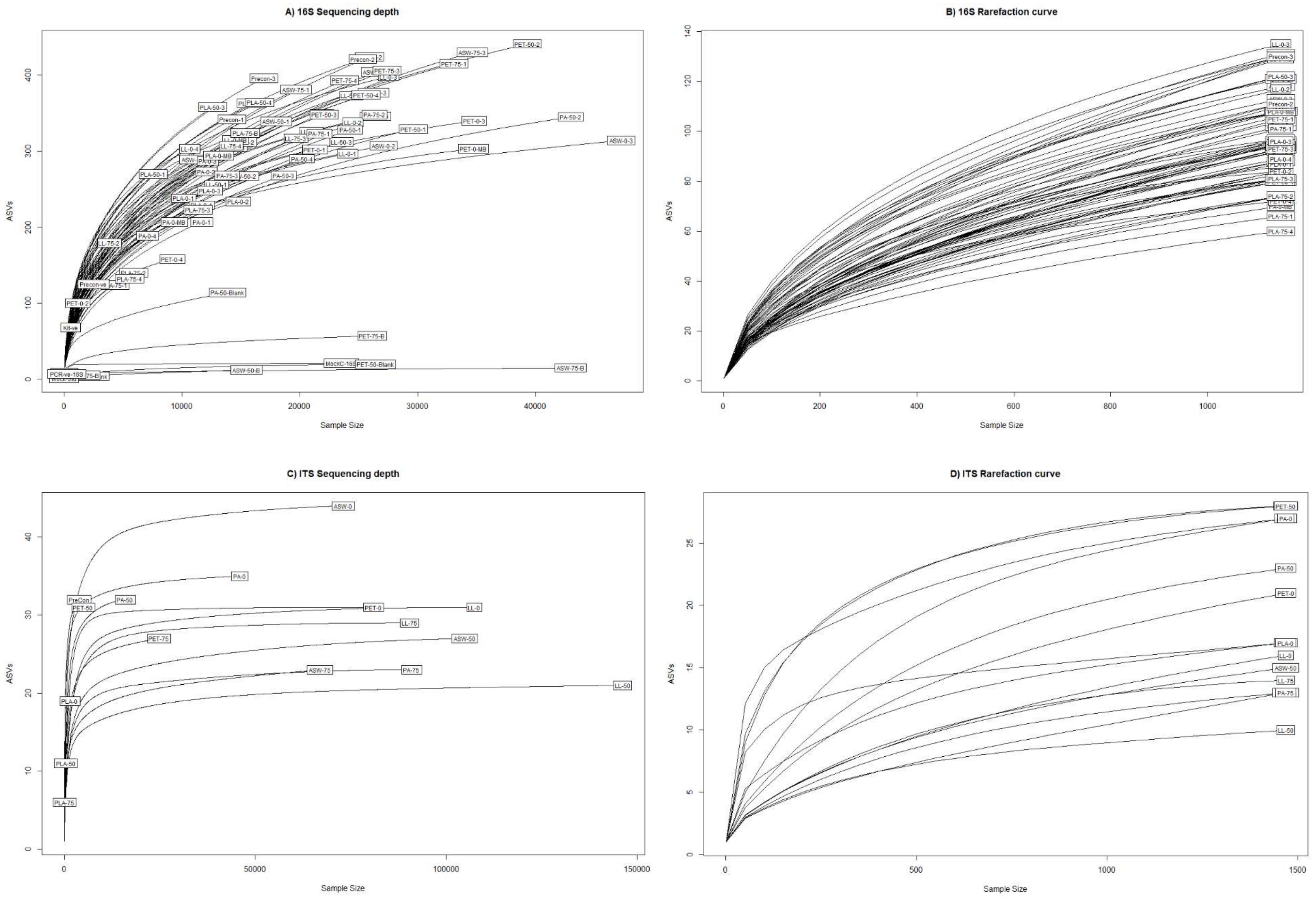


**Figure S9.** Rarefaction curves of ASV data derived from 16S rRNA and ITS2-4 gene sequence region analysis following rarefaction to lowest sample read number with the exclusion of control negatives. Produced in R version 4.2.2 (R core team, 2020).

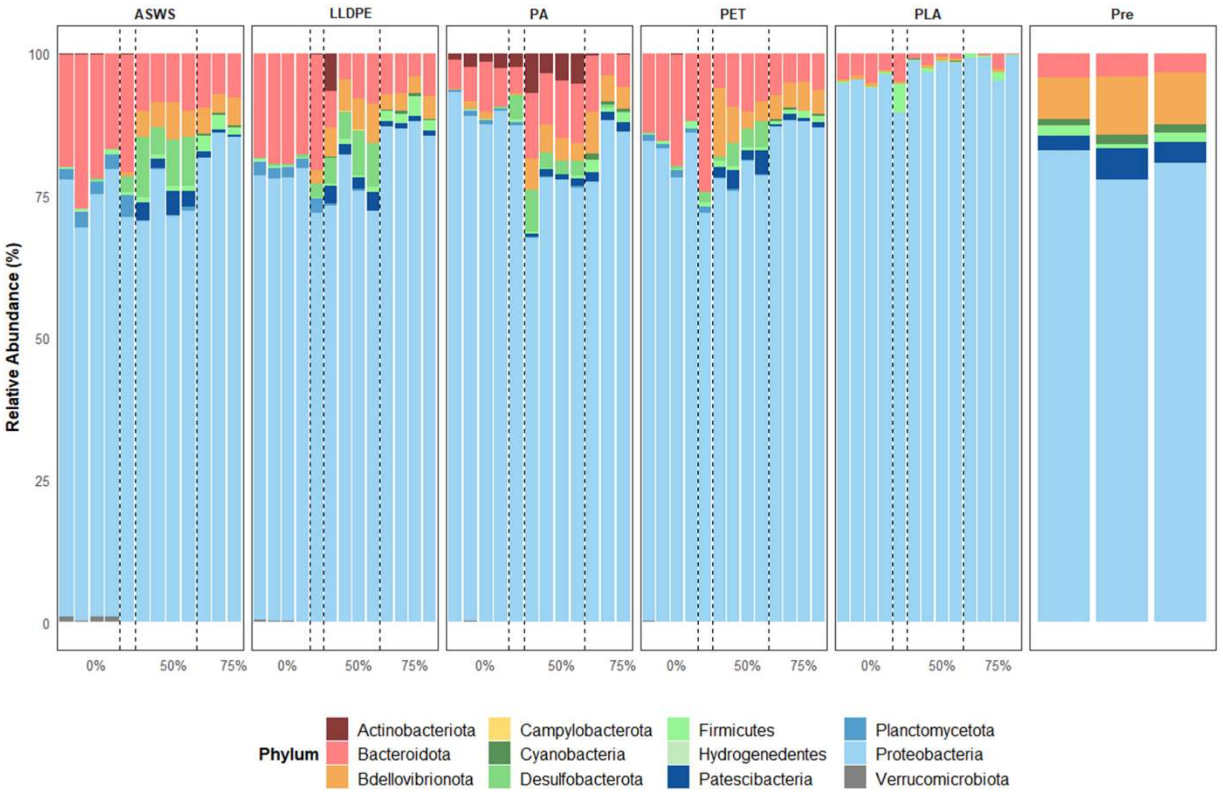


**Figure S10.** Bacterial phyla relative abundance plots, based on analysis of 16S rRNA genes. Samples include (ASWS) artificial seawater broth, (LLDPE) linear low-density polyethylene, (PA) polyamide-6, (PET) polyethylene terephthalate, (PLA) polylactic acid, and (Pre) precondition growth media [Marine Broth 2216]. Following filtering and rarefaction, sequence reads were identified using the DADA2 package in R (R Core Team 2020) for read processing, and a non-redundant SILVA_v138.1 database (McLaren 2020) used for taxonomic assignment for samples which contained at least 1000 sequence reads.

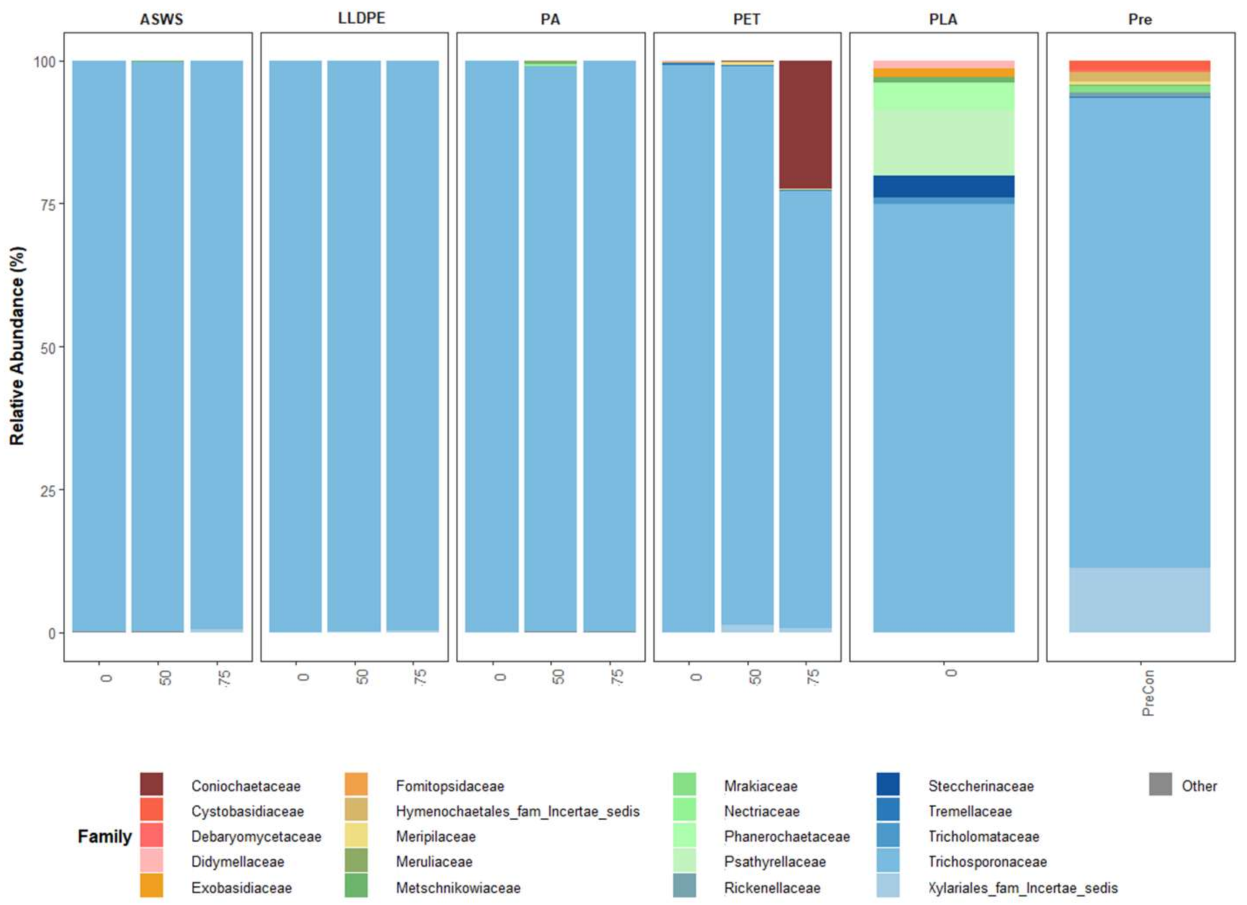


**Figure S11.** Relative abundance of the top 20 most abundant fungal families found in replicate pooled, plastic leachate samples from three different growth conditions: (0) 0% additional carbon, (50) 50% residual carbon supplemented with Marine broth 2216 media (75) 75% supplemented carbon with Marine Broth 2216 media. Samples include (ASWS) artificial seawater broth, (LLDPE) linear low-density polyethylene, (PA) polyamide-6, (PET) polyethylene terephthalate, (PLA) polylactic acid, and (Pre) precondition growth media [Marine Broth 2216]. Samples containing less than 1000 sequence reads (PLA-50 and PLA-75) were removed during filtering.

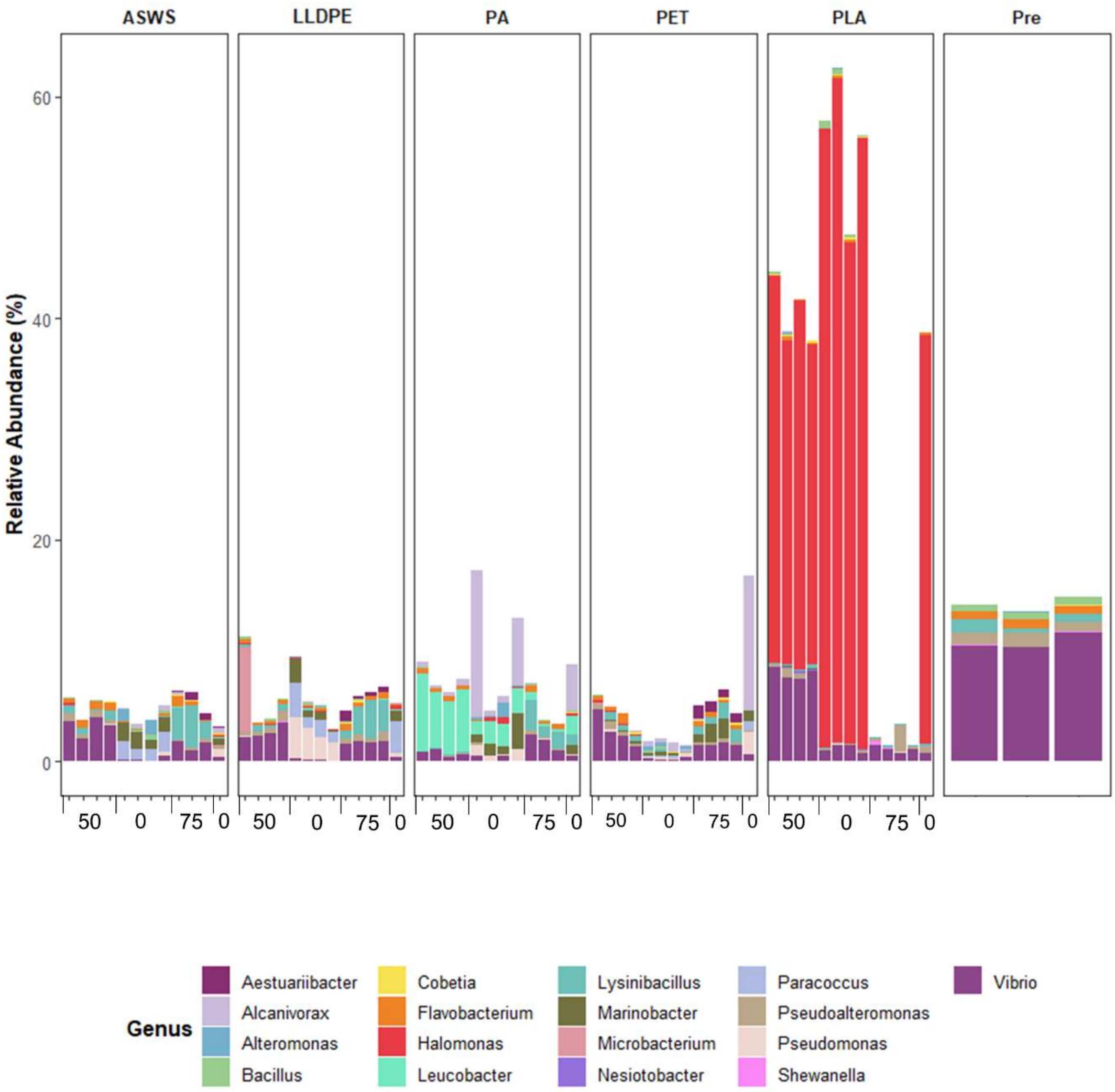


**Figure S12.** The relative abundance of previously reported putative plastic bacterial degraders as described in a microbial plastic degraders database (https://plasticdb.org). Samples included artificial seawater broth (ASWS) and media containing 'leachate' from linear low-density polyethylene (LLDPE), polyamide-6 (PA), polyethylene terephthalate (PET), polylactic acid (PLA), and preconditioned growth media [Marine Broth 2216] (Pre).

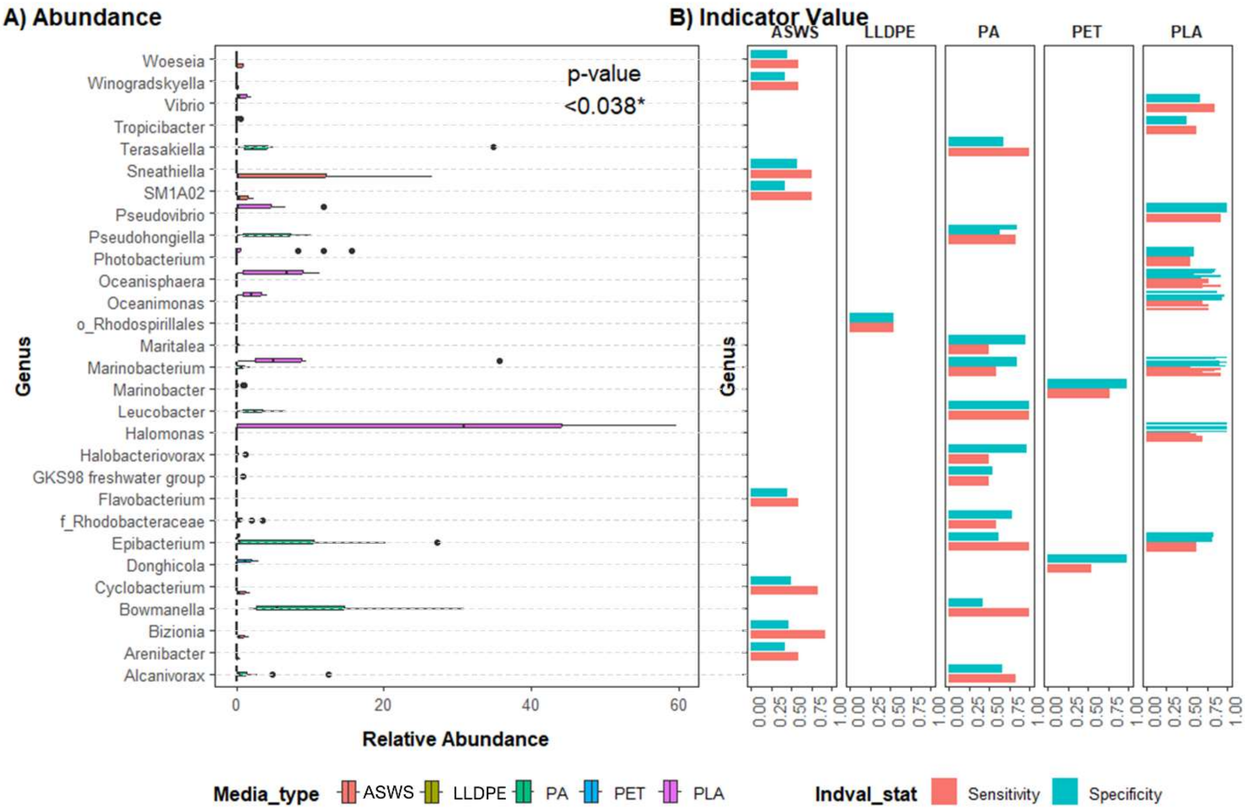


**Figure S13.** A) Box and whisker relative abundance of genera identified as having statistically significant associations with a plastic leachate type. Box mean lines indicate the average abundances of ASV assigned to each genera, where whiskers highlight the upper and lower quartile of ASV abundances, and dots signify outlier abundance. These genera are considered indicator species for the respective plastic leachate types based on default parameters of a multi-level patterns analysis. All taxa had a p-value $\leq 0.04$. B) Indicator specificity and sensitivity values result from multi-level pattern analysis. Specificity refers to the presence of the taxa within the media, and sensitivity refers to its abundance. Treatments consisted of an artificial seawater broth and four plastic leachate types: linear low-density polyethylene (LLDPE), polyamide-6 (PA), polyethylene terephthalate (PET) and polylactic acid (PLA).

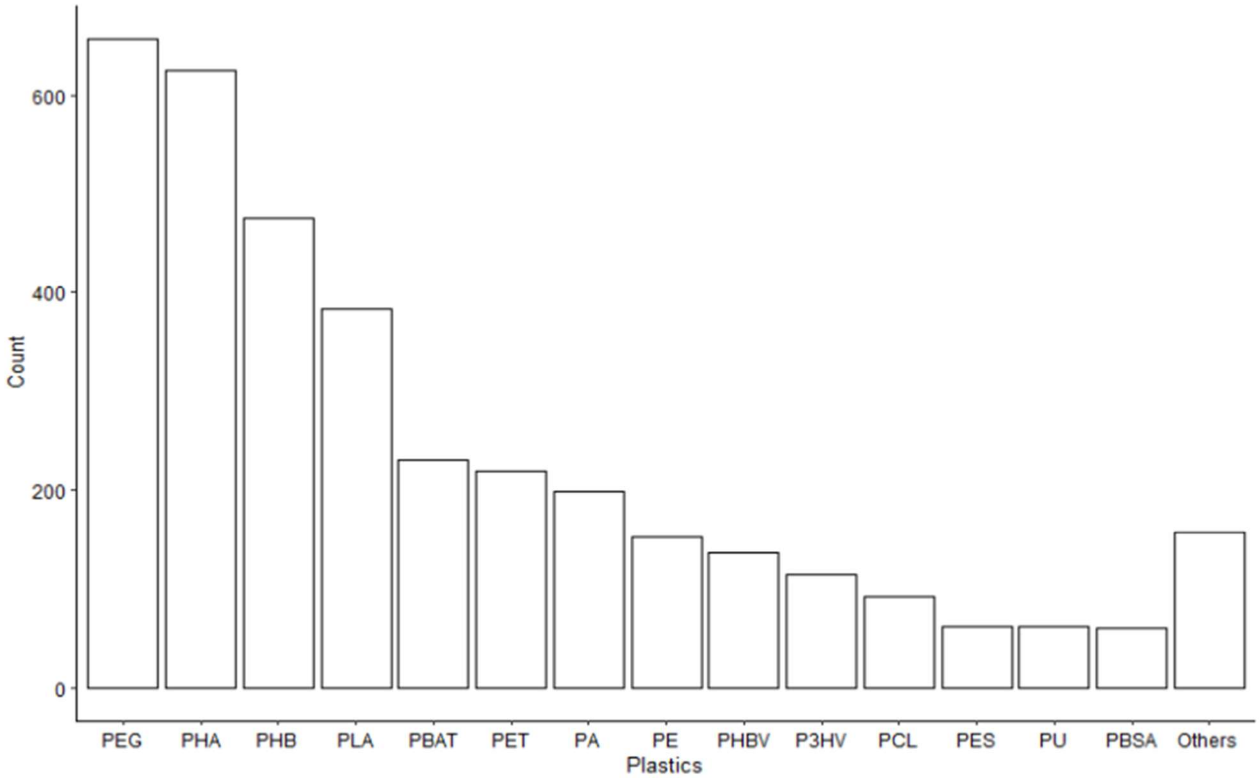


**Figure S14**. Total number transcripts identified within the assembled meta-transcriptome matching genes reported as coding for plastic degrading enzymes as characterized by PlasticDB (Gambarini et al. 2022), with a minimum e-value of 1.0 $e^{-06}$ and percentage identity match of 30 %.

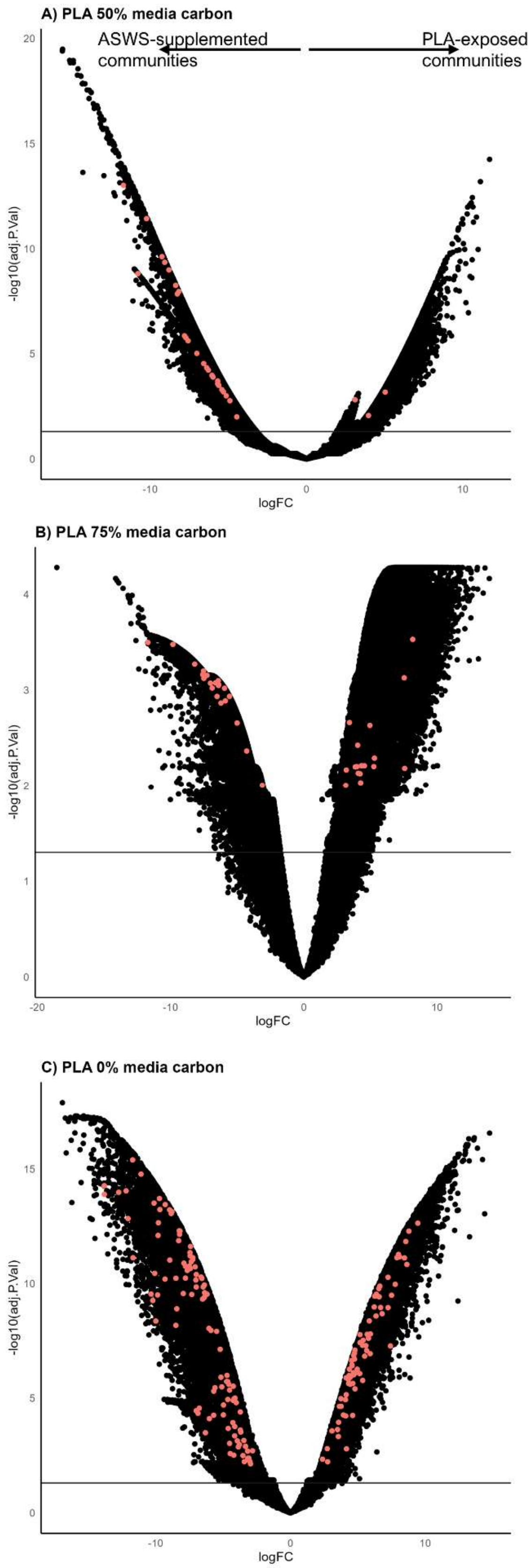


**Figure S15.** Differentially expressed transcripts of PLA-leachate-exposed and ASWS-exposed microbial communities based on log-scaled normality of the Limma-voom package in R (Law et al. 2014). Positive log fold change indicates increased abundance in (A) PLA 50% and (B) 75% media carbon transcripts, compared to ASWS 50% and 75% media carbon transcripts. (C) Compares PLA 0% media carbon with the 100% media carbon preconditioned communities. Dots above the horizontal line indicate FDR (false discovery rate) adjusted p-values < 0.05. Coloured dots represent plastic degrading genes as identified by PlasticDB (Gambarini et al. 2022), that were present in at least two of the DEA tools tests (DESeq2 (Love et al. 2014), EdgeR (Robinson et al. 2010) and Limma + voom (Law et al. 2014)).

### Supplementary Tables

**Table S1.** Concentrations of additives and inorganic material in each of the plastic polymer types used in the current study.

| Plastic Type | Base Polymer | Known Additive | *Additive Content | ‡Inorganic content |
|---|---|---|---|---|
| **Linear low-density polyethylene (LLDPE)** | Innoplus LL7420A | Irganox B215 | 0.25% | <0.1% |
| **Polyamide 6 (PA6)** | Ultramid B3S | Nylostab S-EED | 0.50% | <0.1% |
| **Polyethylene terephthalate (PET)** | PAPET COOL IV 0.80 | Tinuvin 234 | 0.30% | <0.1% |
| **Polylactic acid (PLA)** | Ingeo Biopolymer 3025D | No additive | -- | <0.1% |

*Additive contents (%) were provided by the plastic manufacturers or their respective distributors.

‡We calculated the inorganic content of each plastic using thermogravimetric analysis.

**Table S2** Primer pairs used to amplify bacterial and fungal DNA from all extracts. Underlined sequences represent the Illumina Nextera adaptor overhangs required for sample indexing (Kozich et al. 2013).

| Target | Forward Primer | Reverse Primer | Reference |
|---|---|---|---|
| [1]**Bacteria (*16S rRNA* gene)** | TCG TCG GCA GCG TCA GAT GTG TAT AAG AGA CAG CCT ACG GGN GGC WGC AG | GTC TCG TGG GCT CGG AGA TGT GTA TAA GAG ACA GGA CTA CHV GGG TAT CTA ATC C | (Klindworth et al. 2013) |
| [2]**Fungi (*ITS2* region)** | TCG TCG GCA GCG TCA GAT GTG TAT AAG AGA CAG GTG ART CAT CGA ATC TTT G | GTC TCG TGG GCT CGG AGA TGT GTA TAA GAG ACA GTC CTC CGC TTA TTG ATA TGC | (Ihrmark et al. 2012) |

[1]Bacterial PCR protocol parameters: Initial denaturation (95°C for 3 min); 25 cycles of denaturation (95°C for 30 sec), annealing (55°C for 30 sec), extension (72°C for 30 sec); final extension (72°C for 5 min).

[2]Fungal PCR protocol parameters: Initial denaturation (94°C for 5 min); 30 cycles of denaturation (94°C for 30 sec), annealing
